# Spatially Resolved Transcriptomic Signatures of Hippocampal Subregions and *Arc*-Expressing Ensembles in Active Place Avoidance Memory

**DOI:** 10.1101/2023.12.30.573225

**Authors:** Isaac Vingan, Shwetha Phatarpekar, Victoria Sook Keng Tung, A. Iván Hernández, Oleg V. Evgrafov, Juan Marcos Alarcon

**Author notes:** **Correspondence:** Juan Marcos Alarcon.

## Abstract

The rodent hippocampus is a spatially organized neuronal network that supports the formation of spatial and episodic memories. We conducted bulk RNA sequencing and spatial transcriptomics experiments to measure gene expression changes in the dorsal hippocampus following the recall of active place avoidance (APA) memory. Through bulk RNA sequencing, we examined the gene expression changes following memory recall across the functionally distinct subregions of the dorsal hippocampus. We found that recall induced differentially expressed genes (DEGs) in the CA1 and CA3 hippocampal subregions were enriched with genes involved in synaptic transmission and synaptic plasticity, while DEGs in the dentate gyrus (DG) were enriched with genes involved in energy balance and ribosomal function. Through spatial transcriptomics, we examined gene expression changes following memory recall across an array of spots encompassing putative memory-associated neuronal ensembles marked by the expression of the IEGs *Arc*, *Egr1*, and *c-Jun*. Within samples from both trained and untrained mice, the subpopulations of spatial transcriptomic spots marked by these IEGs were transcriptomically and spatially distinct from one another. DEGs detected between *Arc*+ and *Arc*-spots exclusively in the trained mouse were enriched in several memory-related gene ontology terms, including “regulation of synaptic plasticity” and “memory.” Our results suggest that APA memory recall is supported by regionalized transcriptomic profiles separating the CA1 and CA3 from the DG, transcriptionally and spatially distinct IEG expressing spatial transcriptomic spots, and biological processes related to synaptic plasticity as a defining the difference between *Arc*+ and *Arc*-spatial transcriptomic spots.

## 1 INTRODUCTION

The neural operations supporting memory require the involvement of various brain systems interconnected through neural networks (Kitamura et al., 2017). The activity of these networks is fine-tuned by experience through the selective recruitment of ensembles of neurons ascribed to contain the bits of information associated with memory (Josselyn and Tonegawa, 2020). Neuronal ensemble activity is shaped by synaptic plasticity mechanisms that modulate the weight and efficacy of the ensemble’s synaptic connections (Takeuchi et al., 2014, Malenka and Bear, 2004). Changes in gene expression are a key underlying mechanism in synaptic plasticity (Mayford et al., 2012), and recent studies have identified multiple profiles of gene expression associated with memory (Marco et al., 2020, Rao-Ruiz et al., 2019, Reshetnikov et al., 2020).

In the rodent hippocampus, encoding of spatial memory information is supported by the diversity of synaptic computations within and across its sub-fields (Poo et al., 2016, Mayford, 2014). Each hippocampal subregion (i.e. Dentate Gyrus, CA1, CA2, CA3) shows distinct patterns of activity to process spatial information and to form or retrieve memories (Middleton and McHugh, 2020, Chevaleyre and Siegelbaum, 2010, Senzai, 2019, Basu and Siegelbaum, 2015, Amaral and Witter, 1989, Sloviter and Lomo, 2012). At the regional level, patterns of activity are conferred through the combination of cyto-architecture, local synaptic circuitry, neuronal cell types and synaptic inputs unique to each region (van Strien et al., 2009, Senzai, 2019, Kesner et al., 2004, Lisman, 1999, Ryan et al., 2015, Hunsaker and Kesner, 2013, Molitor et al., 2021, Rao-Ruiz et al., 2019, Shpokayte et al., 2022, Sullivan et al., 2021, Shah et al., 2016, Cho et al., 2016). At the single-cell level, the functional state of hippocampal neurons is defined by connectivity, hippocampal subregional location, and experience-dependent recruitment into a memory ensemble (Josselyn and Tonegawa, 2020, Tonegawa et al., 2018).

The investigation of memory across molecular and functional levels necessitates an approach that connects the scales of hippocampal organization between cellular mechanisms and networks of cells. Though hundreds of individual genes supporting synaptic plasticity at the cellular level have been identified (Sanes and Lichtman, 1999, Alberini and Kandel, 2014, Asok et al., 2019, Magee and Grienberger, 2020, Malenka and Bear, 2004), the transcriptomic profiles which characterize memory across hippocampal subregions and memory-associated neuronal ensembles are understudied.

Memory-associated neuronal ensembles are a sparsely distributed population of neurons which are strongly activated together during the acquisition and recall of a memory, and are thought to encode the traces of information relevant to that memory (Ryan et al., 2015, Guan et al., 2016, Chen et al., 2019). Numerous studies have identified Immediate Early Genes (IEGs) such as *Arc*, *c-Fos*, *Egr1*, *c-Jun*, *Npas4*, *Bdnf*, and *Fmrp* to characterize memory-associated neuronal ensembles (Sauvage et al., 2019, Josselyn and Tonegawa, 2020, Meenakshi et al., 2021, Erwin et al., 2020). In this study, the location of memory-associated neuronal ensembles is inferred from the expression of IEGs in spatial transcriptomic spots. The expression of *Arc* mRNA has been used to tag and characterize the properties of memory-associated neuronal ensembles (Guzowski et al., 1999, Ramirez-Amaya et al., 2005, Lee et al., 2022). This proxy for the memory-associated neuronal ensemble was used to explore the transcriptomic changes supporting memory recall in the hippocampus. As such, we characterized changes in gene expression across the major subregions of the transverse dorsal hippocampus using bulk RNA sequencing of micro-dissected dorsal hippocampal subregions and investigated the spatial distribution of IEG-associated gene expression through a spatial transcriptomic study of coronal dorsal hippocampal slices.

## 2 METHODS

### 2.1 Animals

A total of 18 adult ArcCreERT2::eYFPflx mice (Denny et al., 2014) with C57BL/6 genetic background aged 3-4 months were utilized across all experimental cohorts (Bulk RNA sequencing – 3 male and 3 female, spatial transcriptomics – 2 male, RT-qPCR – 10 male). Prior to experimental onset mice were bred in-house at the SUNY Downstate Health Sciences University vivarium (Brooklyn, NY, USA). Mice were housed in groups of two to five per cage. Beginning on the first day of training and testing, mice were single housed in shoebox cages in a sound attenuation cubicle (Med Associates). For every experimental run, a minimum of two mice —each undergoing the same behavioral conditioning— were simultaneously housed in the sound attenuation cubicle. Ad libitum food and water was provided. Mice were randomly assigned to behavioral cohorts before the start of the experiment. Mice were handled daily for 3 days prior to the start of the experiment to reduce anxiety and improve voluntary approach. All animal procedures proposed are approved by, and will be performed following, the Institutional Animal Care and Use Committee guidelines at SUNY Downstate Health Sciences University.

### 2.2 Behavior

All procedures were performed in compliance with the Institutional Animal Care and Use Committee of the State University of New York, Downstate Health Sciences University. ArcCreERT2::eYFPflx male mice were trained in a hippocampus-dependent two-frame active place avoidance task. The place avoidance system consisted of a 40-cm diameter arena with a parallel rod floor that could rotate at 1 rpm. The position of the animal was tracked using PC-based software (Tracker, Bio-Signal Group Corp., Brooklyn, NY) that analyzed 30-Hz digital video images from an overhead camera. Mice in the trained condition learned the “Room+Arena−” task variant. Place avoidance of a 60° zone was reinforced by a constant current foot shock (60 Hz, 500ms, 0.2mA) that was scrambled (5-poles) across pairs of the floor rods. Rotation of the arena would carry the mouse into the shock zone unless the animal actively avoided the zone. Entering the shock zone for more than 500 ms triggered shock. Additional shocks occurred every 1.5 seconds until the animal left the shock zone. Measures of place avoidance were computed by TrackAnalysis software (Bio-Signal Group Corp., Brooklyn, NY).

On day one, mice received a 30-min trial with the shock off to habituate to the rotating arena. Across the next two days the animals experienced four training trials, with two 30-minute trials a day with the activated shock with a 40 min inter-trial-interval. Control (untrained mice) experienced identical training conditions, except for the shock always being off. Memory performance was assessed 24 hr. after the final training session in a 10-minute retention test with the shock off.

### 2.3 Microdissected bulk RNA sequencing

60 minutes post retention test, mice were euthanized, and brains were extracted, washed in ice cold artificial cerebrospinal fluid, blocked, and mounted on a vibratome stage. Hippocampal subregions (DG, CA3, CA1) were microdissected from 400 µm thick coronal live tissue sections. Subregions from the dorsal hippocampus were extracted and microdissected using microsurgical tools. Microdissected pieces of tissue from each animal were pooled by subregion in 500 µL of prechilled TRIzol in a 1.5mL microcentrifuge tubes and stored at -80°C.

Total RNA was extracted using the Direct-zol RNA miniprep kit (Zymo Research, Irvine, CA, United States) according to the manufacturer’s protocol. RNA quality was assessed on an Agilent 2200 TapeStation (Agilent Technologies, Palo Alto, CA, United States). Samples with a RIN greater than 7 were processed for library preparation with NEBnext rRNA depletion kit and ULTRAII FS RNA-seq Library Preparation Kit for Illumina (New England Biolabs, Ipswich, MA, United States). This procedure involved steps for mRNA enrichment, fragmentation, random primed cDNA synthesis, second strand synthesis, end repair, A-tailing, adaptor ligation, and PCR amplification. Size selection and cleanup was performed with SPRI select beads (Beckman Coulter, Indianapolis, IN, United States). Sequencing was performed on the Novaseq6000 (Illumina, Inc., San Diego, CA, United States) to obtain paired end sequencing reads.

### 2.4 Bulk RNA sequencing data preprocessing

RNA sequencing reads obtained from NovaSeq 6000 were systemically processed to ensure high-quality outputs for downstream analyses. To evaluate the quality of the reads, quality control analysis was performed using *FASTQC* v0.11.9. RNA sequencing reads in Binary Base Call (BCL) format were subsequently converted using *bcl2fastq2* v2.20 to FASTQ format, and simultaneously demultiplexing to assign sequences to their respective samples based on index sequences. Next, FASTQ files were then mapped to mouse MM10 (GRCm39) reference genome using *STAR* 2.7.10a aligner. Lastly, the transcript quantification of the reads was done using *Salmon* v1.2.0 to create a count matrix for downstream analyses.

### 2.5 Bulk RNA sequencing analyses

We conducted several differential expression analyses using *DESeq2* v1.1.0 in the Illumina BaseSpace, comparing RNA counts between trained and untrained sample groups. Genes with a false discovery rate (FDR) below 0.05 were deemed differentially expressed. To identify the variances in our count data, we performed principal component analysis (PCA) on the normalized count data, encompassing all hippocampal regions and within each specific region.

To visualize our data, we generated a heatmap in *ComplexHeatmap* v2.18.0 using Z-scores, which standardized the expression levels of genes across samples (Gu et al., 2016). The heatmap, organized with dendrogram and sorted using Euclidean distance, allowed us to identify patterns of similarity in gene expression across samples.

To explore deeper into biological significance, we conducted Gene Ontology (GO) enrichment analysis using *clusterProfiler* v4.10.0 in R (Wu et al., 2021), focusing on biological processes, cellular components, and molecular functions. Only enriched GO terms with an FDR less than 0.01 were identified as significant, and the top 10 GO terms were displayed in a ranked dot plot. Redundant terms were filtered out using *simplify* with a cutoff of 0.7. The top 10 enriched pathways per category were visualized in dot plots, ordered by significance.

We also utilized *Vennplex* v1.0.0.2 software to compare and visualize the overlaps among differentially expressed genes (DEGs) from our differential expression analyses (Cai et al., 2013). The gene lists were categorized into upregulated and downregulated, allowing the visualizing of the gene overlaps and identification of counter-regulated genes (genes exhibiting opposite expression trends in different regions). We employed the *SuperExactTest* v1.1.0 in R to statistically assess whether the observed overlapping genes across different hippocampal regions significantly exceeded the expected overlap by random coincidence (Wang et al., 2015). The assessment was performed using hypergeometric distribution and Fisher’s Exact test, and the statistical significance of these gene overlaps was shown in upset-style plots.

### 2.6 Tissue preparation for spatial transcriptomics

60 minutes after retention test, mice were euthanized and brains were extracted and immediately prepared for snap freezing in cuvettes of Tissue-Tek Optimal Cutting Temperature compound (Sakura Finetek USA, Torrance, CA, United States) floating in a bath of methyl-butane chilled by liquid nitrogen. Brain blocks were cryosectioned at -20°C (10x Genomics, 2023) and 10 µm thick coronal sections containing the dorsal portion of the hippocampus were obtained (bregma - 1.7 and -2.2 mm (Paxinos and Franklin, 2019)). Collected tissue sections are trimmed to fit within the 5 mm x 5 mm capture area on the Visium gene expression slide. Sections were mounted on a prechilled Visium gene expression slide or a Tissue Optimization slide.

For Optimization, all tissue sections were collected from a single tissue. For Visium gene expression, slide capture areas contained one section per biological sample. Slides were fixed in prechilled methanol before staining using the Visium spatial tissue optimization protocol (Genomics, 2019) or the Visium spatial gene expression protocol (Genomics, 2023).

Tissues permeabilization time on the gene expression slide was set to 18 minutes based on our tissue optimization results. Images were taken according to the Visium Spatial Gene Expression Imaging Guidelines(Genomics, 2021). Brightfield histology images for the Visium Tissue Optimization and Gene Expression slides were taken using the Leica Aperio CS2 Slide Scanner at 20x magnification. Tissue optimization fluorescent images were taken on a Zeiss LSM 800 (555nm LED,75% intensity, and 200ms exposure).

mRNA was extracted and libraries were prepared following the Visium Spatial Gene Expression User Guide (Genomics, 2023). They were loaded at 300 pM and sequenced on a NovaSeq 6000 System (Illumina) using a NovaSeq S4 Reagent Kit (200 cycles, catalog no. 20027466, Illumina), targeting a sequencing depth of approximately 250–400 10 × 106 read-pairs per sample. Sequencing was performed using the following read protocol: read 1: 28 cycles; i7 index read: 10 cycles; i5 index read: 10 cycles; and read 2: 91 cycles.

### 2.7 Spatial transcriptomics preprocessing

Raw RNA sequencing data obtained from NovaSeq 6000 were systemically processed to ensure high-quality outputs for downstream analyses. Raw sequencing data in BCL format were converted using *bcl2fastq2* v2.20 to FASTQ format, and simultaneously demultiplexing to assign sequences to their respective samples based on index sequences. Following the conversion, these FASTQ files were subsequently processed with *TrimGalore* v.0.6.5 to automate quality, adapter trimming and perform quality control.

For the alignment and the quantification of gene expression, *Space Ranger* v2.0.0 (10x Genomics) was used to generate and quantify raw gene-barcode matrices from RNA sequencing data. Alignment and quantification of Visium spatial gene expression data utilizes the same tools implemented in the analysis of single cell RNA-seq data sets. Data were aligned to refdata-gex-mm10-2020-A, a mouse genome index provided by 10x Genomics, which is an annotated version of mm10 genome assembly. The gene-barcode matrices identified the number of unique molecular identifiers (UMIs) associated with each gene for each cellular barcode, allowing the quantification of transcript abundance.

After the alignment and quantification, Space Ranger toolset was utilized to normalize the gene expression data within the raw gene-barcode matrices to account for any technical variations. Finally, we also utilized Space Ranger to align gene expression data with spatial coordinates derived from histological images of each sample, facilitating the visualization of transcriptional activity to specific tissue location.

### 2.8 Spatial transcriptomics analysis

Gene expression outputs from Space Ranger were analyzed using the software package *Seurat* v5 in the R coding environment (Hao et al., 2023, Butler et al., 2018). Spatial gene expression data were cropped to only include the capture spots within the dorsal hippocampus. Using default Seurat commands, data from each sample were normalized, transformed (using *SCTransform v2)* and integrated. The integrated data set were further normalized and clustered (resolution = 0.8) in PCA space. Clustered data can be visualized through UMAP projections of the capture spots or in the 2D space of the tissue section. Cell type annotation was performed by integrating our data with a random subset of 10,000 hippocampal cells from the Allen Brain Atlas Cortex and Hippocampus Single Cell Taxonomy (Yao et al., 2021). Predicted cell-type identities were calculated and optimized based on gene expression similarity and anatomical location.

Capture spots were grouped along two different criteria to calculate differential gene expression in our samples. First, capture spots belonging to the whole hippocampus and each anatomical cell layer in the DG, CA3 and CA1 regions were compared across training conditions. Next, within each training condition, hippocampal capture spots were separated into two groups based on the detectable (>0) expression of the immediate early gene (e.g. *Arc+* and *Arc-* groups). IEG expressing and not expressing groups of spots were compared both within training conditions and across training conditions. Differential gene expression was calculated using the Wilcoxon rank sum test (the default method for Seurat). Genes with an of FDR below 0.05 were considered differentially expressed genes/transcripts. GO term enrichment and overlap analyses on detected DEGs were performed as described above for bulk RNA sequencing DEGs.

Expression data for 23 IEGs from all hippocampal spots evaluated through pairwise Pearson’s Correlations using the R-language base stats package. Pearson’s correlation coefficients between all IEG pairs were calculated within samples, alongside P values to assess significance. Heatmaps of the correlation matrix were plotted using the R-language package, ComplexHeatmap.

### 2.9 RT-qPCR

Dorsal hippocampal subregions (CA1, CA3, and DG) were microdissected as described for bulk sequencing. Tissue was homogenized using TRIzol (Thermo Fisher Scientific, Waltham, MA, United States) before RNA extraction by guanidinium-phenol-chloroform extraction with ethanol precipitation was performed and quantified using spectrophotometry. cDNA was reverse transcribed from sample bulk RNA using Thermo Fisher’s SuperScript IV Kit (Thermo Fisher Scientific, Waltham, MA, United States). Relative gene expression was quantified using the SYBR green master mix system (Ponchel et al., 2003). 10 µL reactions were prepared in the wells of a 384 well plate, hippocampal regions were each prepared on separated plates. Each well contained 2.5ng of template cDNA and 0.5 µM each of forward and reverse primers for one gene per well for a total reaction volume of 10µL (sequences and melting temperatures can be found in table 1). Each gene was loaded in triplicate in each plate to account for pipetting errors. Plates were incubated in the Bio-Rad CFX384 Real-Time PCR Detection System (Bio-Rad Life Sciences, Hercules, CA, United States) using the following schedule: 50°C for 5 min (1 cycle), 95°C for 10 min (1 cycle), 60°C for 1 min (1 cycle), and 95°C for 15 s followed by 60°C for 1 min (40 cycles).

**Table 1.**
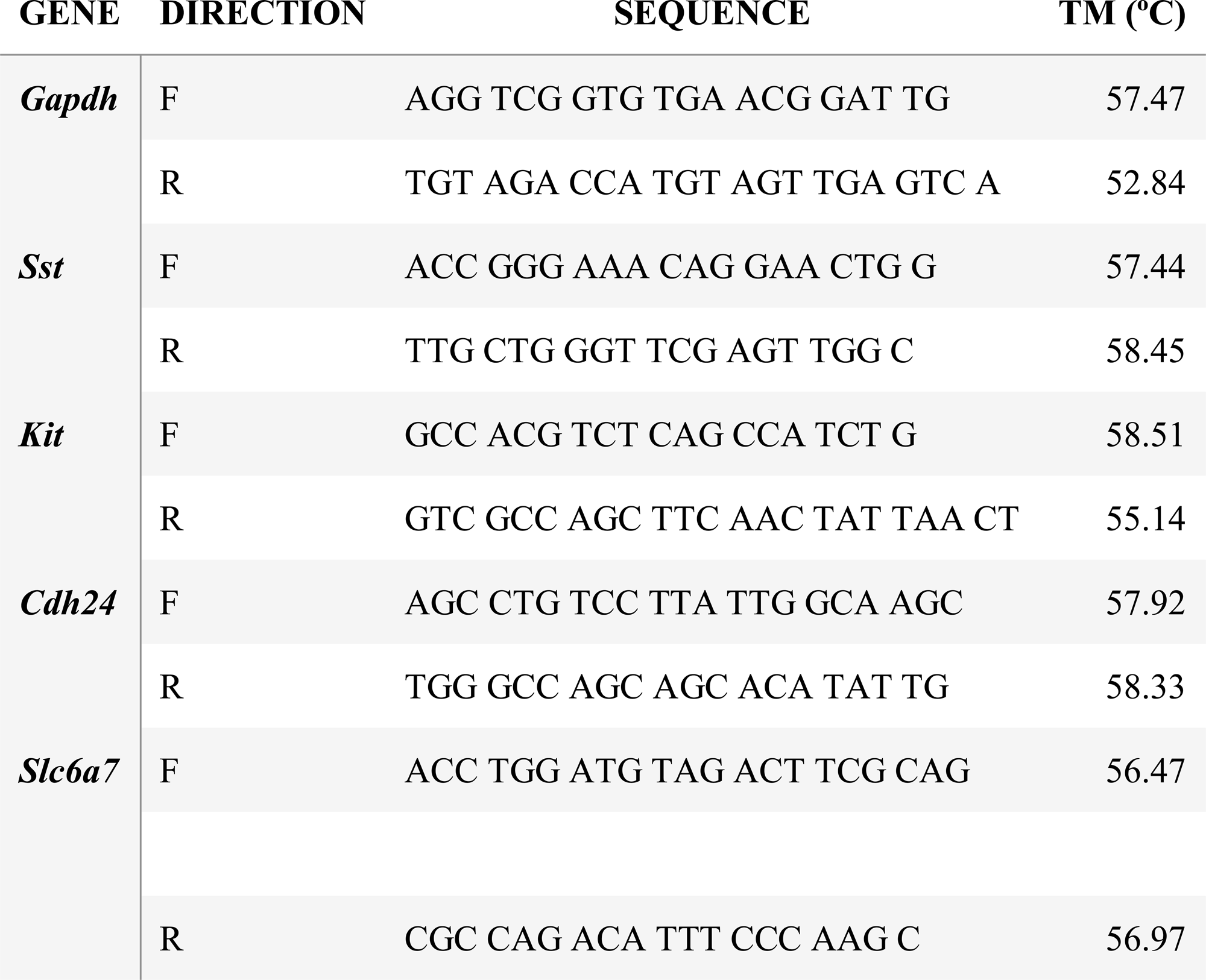
RT-qPCR primer sequences.

Ct values for each reaction are automatically determined (roughly 20% of the plateau value). Data were analyzed for relative gene expression quantification utilizing the ΔΔCt method, using the expression of *Gapdh* as the internal control. Relative expression values were normalized to the expression value of the untrained group and Student’s t-tests were performed on each primer target across training conditions using the ΔCt value for each comparison. Benjamini-Hochberg correction was performed to adjust p-values for multiple hypothesis testing.

## 3 RESULTS

### 3.1 APA trained mice exhibit place avoidance memory

To investigate gene expression changes induced by active place avoidance (APA) training in the hippocampal network, we conducted bulk RNA-sequencing and spatial transcriptomics (see Figure 1A for experimental timeline). In the APA, mice learn to avoid a 60^◦^ shock zone on a rotating circular arena (Cimadevilla et al., 2000, Stuchlík et al., 2013). Mice trained in the APA exhibited a decrease in the number of shocks delivered during the acquisition trials (Figure 1B) and longer re-entry times to the shock zone during the memory retention test trials (Figure 1C).

**Figure 1.**
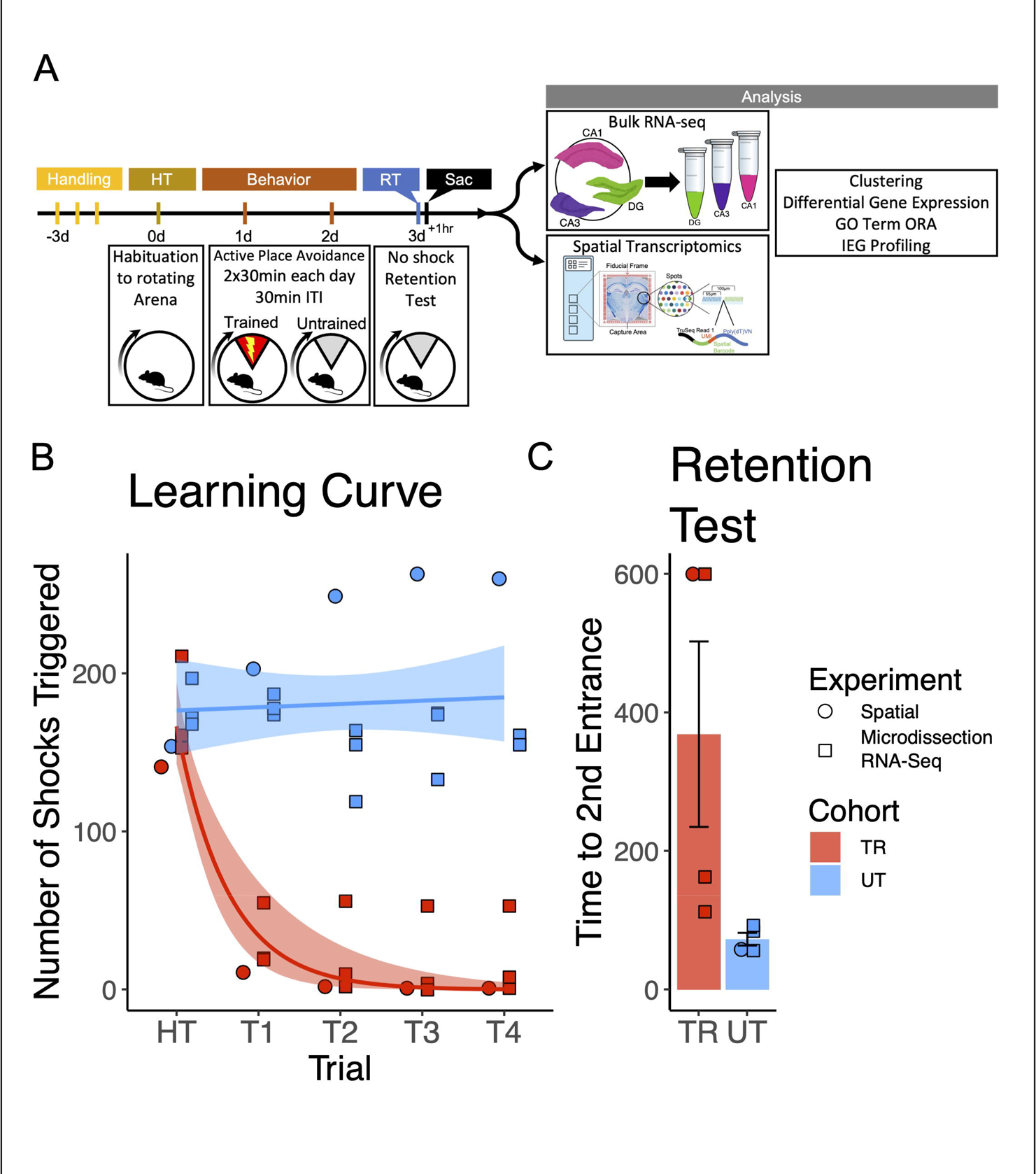
Trained mice learned to avoid the location of an unmarked shock zone. **(A)** Experimental timeline. Mice were trained in the active place avoidance paradigm over the course of 4 days following 3 days of handling. On day 0 they were habituated to a rotating arena with no active shock zone (HT). On day 1 and 2 trained mice received two 30-minute trials with an active shock zone. Untrained mice were not exposed to an active shock zone. On day 3 all mice received a 10-minute retention test trial (exposed to the rotating arena with no active shock zone). Mice were sacrificed 60 minutes following the end of the retention test, brains were collected for further processing. **(B)** Training performance measured as number of shocks triggered. Trained mice learned to avoid the location of the shock zone indicated by the decreased number of shocked triggered. Exponential fit learning curves included with shaded error bars (SE). **(C)** Memory performance measured as time to second entrance in the retention test trial. Trained mice demonstrated higher time to second entrance during the retention test. Trained mean 375s ± 70.4 (SE), Untrained mean 58.7s ± 8.51 (SE).

### 3.2 Hippocampal subregional transcriptomes are stratified by training condition

To explore the molecular mechanisms underlying memory recall across the dorsal hippocampus, bulk RNA-sequencing was performed on the Dentate Gyrus (DG), CA3, and CA1 hippocampal subregions. Principal component analysis (PCA) from all 3 subregions combined reveals a separation of samples by behavioral cohort (Figure 2A), indicating that differences in training contribute substantially to the overall variability of gene expression in our samples. Separation between samples is further accentuated when PCA is performed separately on data from each subregion (Figure 2C-D). Consistently, hierarchical clustering of gene expression profiles reliably grouped samples by their behavioral cohort (Figure 2E).

**Figure 2.**
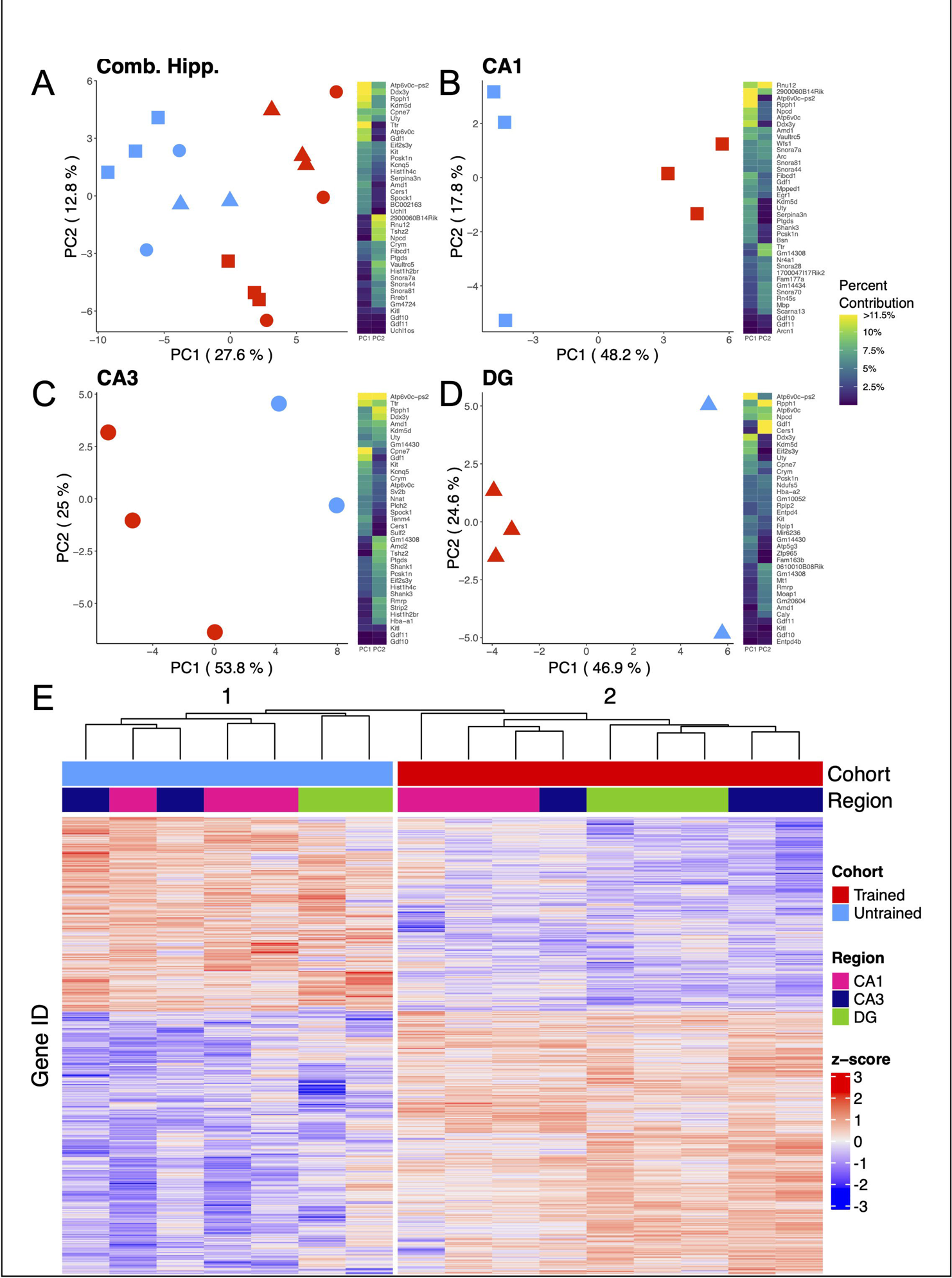
APA training results in substantial changes in hippocampal subregion gene expression profiles. **(A-D)** Left, principal component analyses of all hippocampal subregions **(A)**, CA1 **(B)**, CA3 **(C)**, DG **(D)** illustrate a clear separation between samples collected from trained and untrained mice along the axis that explains the most variance. Right, heatmap with top 20 contributing genes to the first two principal components of each are reported in the heat map to the right of each plot. Sample size for the CA1 was 3 trained and 3 untrained. Sample size for the CA3 was 3 trained and 2 untrained. And Sample size for the DG was 3 trained and 2 untrained. **(E)** Normalized gene expression of the top 2000 most significant DEGs. Hierarchical clustering of gene expression profiles plotted as a dendrogram clusters samples by training condition.

### 3.3 Hippocampal subregional enrichment of learning and memory-related biological processes

We detected 1663, 606, 315 and 350 differentially expressed genes (DEGs) (adjusted p-value < 0.05) in the combined hippocampal subfields, CA1, CA3 and DG respectively, when contrasting APA-trained and untrained mice. (Figure 3A-D & Supplemental Figure 1A-D). Despite high statistical significance (FDR<0.005), most DEG in each comparison exhibited modest fold change (|log_2_ fold| < 1.0) difference between the experimental group. Next, we investigated the overlap of regionally detected DEGs (FDR < 0.05) to elucidate the distribution of DEGs across hippocampal subregions. We found the largest overlap of upregulated DEGs between the CA3 and CA1 subregions with a total of 127 intersecting genes (Figure 3E & Supplemental Figure 1E).

**Figure 3.**
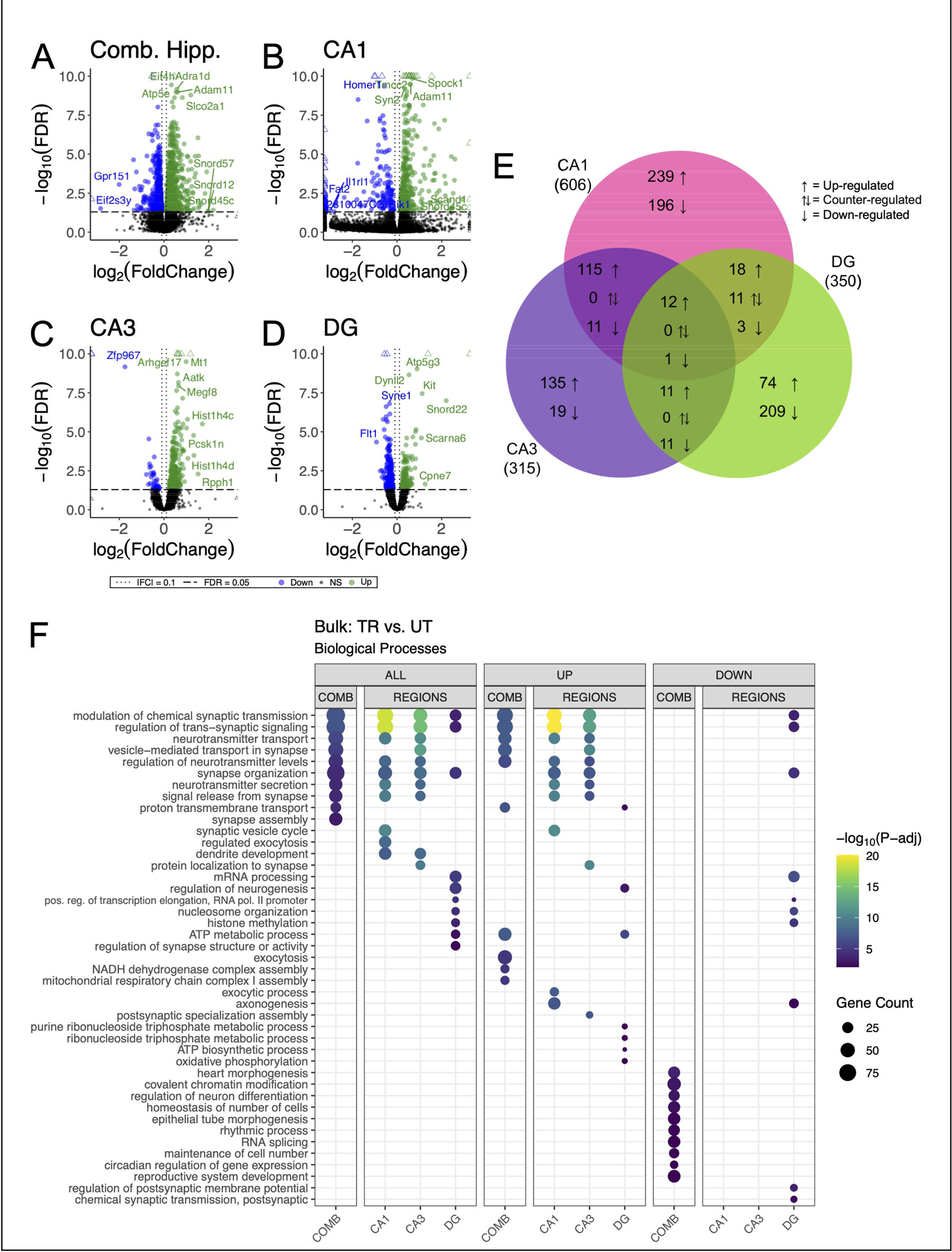
Bulk RNA-seq of hippocampal subregions reveals similar expression profiles in CA1 and CA3 regions in trained samples. **(A-D)** Volcano plot of DEGs between trained and untrained animals in all hippocampal subregions combined **(A)**, and in the CA1 **(B)**, CA3 **(C)**, and DG **(D)** subregions individually. In all hippocampal subregions combined, 966 genes were upregulated and 697 were downregulated. 393 genes were upregulated and 213 were downregulated in the CA1, 273 genes were upregulated and 42 were downregulated in the CA3, and 117 genes were upregulated and 223 were downregulated in the DG. See Supplemental Figure 1A-D for full sized plots. **(E)** Overlaps in regional DEGs demonstrate greatest similarity between CA1 and CA3 subregions. DEGs from each regional analysis were stratified by direction of fold change. Overlaps in DEGs between the CA3 and CA1 subregions were significant (Fisher’s Exact Test). See Supplemental Figure 1E for further details. **(F)** Regional enrichment of biological processes detected amongst all DEGs (left) and stratified by up-(middle) and down-regulated (right) DEGs. Dot color reflects the statistical significance (-log_10_(FDR)) of the biological process enrichment. Dot size reflects the number of detected DEGs mapped to the genes involved in a given biological process.

GO enrichment analysis was conducted to identify overrepresented biological process among DEGs identified within each hippocampal subregion (The Gene Ontology, 2019). Among the DEGs detected in the combined hippocampal subregions, we found overrepresentation of genes involved in the modulation of neurotransmitter signaling (Figure 3F). These DEGs included *Ywhag*, *Bsn*, and *Nup153*, which have been previously implicated in these cellular functions (Wachi et al., 2017, Annamneedi et al., 2021, Leone et al., 2019). DEGs detected in both the CA3 and CA1 were also enriched with genes involved in the modulation of neurotransmitter signaling. The genes *Snap25, Shank1, Shank 3, Bsn and Ywhag* were identified in the overlap of upregulated DEGs between these regions and have been studied for their involvement in synaptic transmission (Irfan et al., 2019, Mao et al., 2015, Shi et al., 2017, Kouser et al., 2013). DEGs detected in the DG were enriched with genes involved in neurogenesis transcriptional regulation. Among these DEGs were *Kit*, *H2afz*, and *Kmt2e*, which have been studied for their relationship to neurogenesis and transcriptional regulation (Katafuchi et al., 2000, Ferreira et al., 2016, Barbieri et al., 2018, Zovkic et al., 2014, Collins et al., 2019). When enrichment analyses were separated by up- and down-regulated DEGs the differences separating the CA1 and CA3 from the DG became more striking with no significant enrichment of biological processes among the downregulated DEGs of the CA1 and CA3. These data reveal a spatial distribution of gene expression across the dorsal hippocampus, defined by a greater degree of similarity between the CA1 and CA3 than between either of these subregions and the DG.

### 3.4 Differences in the spatial distribution of hippocampal gene expression between trained and untrained animals

To further investigate the spatial distribution of DEGs associated with APA memory across the hippocampus, we performed spatial transcriptomics on coronal sections containing the dorsal hippocampus from one trained and one untrained mouse. Computational analyses of integrated capture spots in the hippocampal slices from trained and untrained animals revealed distinct clusters (seen in a Uniform Manifold Approximation and Projection (UMAP) plot) which map along anatomical boundaries in a spatial transcriptomic plot (Figure 4A, B). Spatial transcriptomic data were integrated with the Allen Brain Atlas (Yao et al., 2021) to computationally annotate each spot with the cell-type most prominently detected (Figure 4C, D).

**Figure 4.**
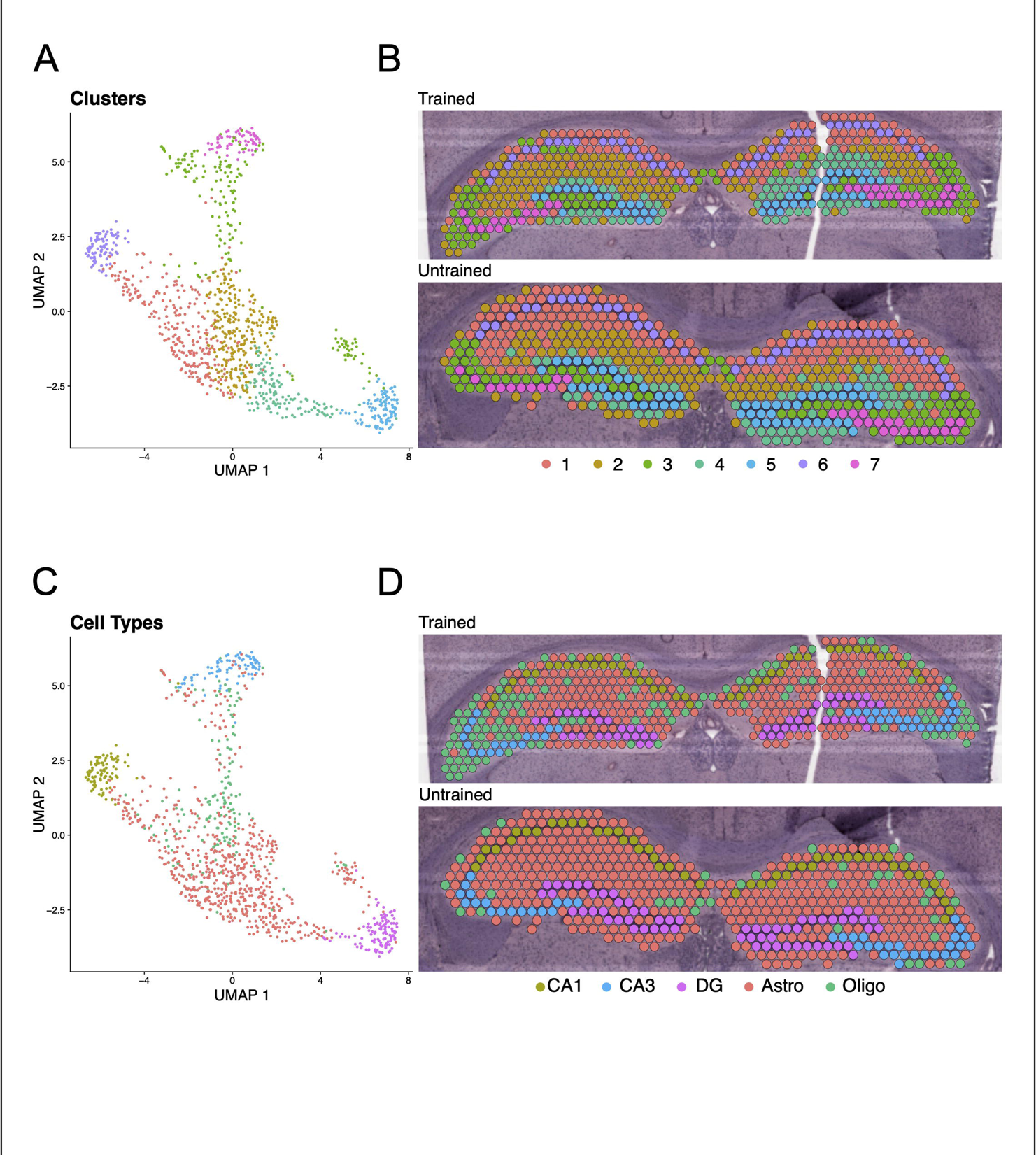
Clustering and cell-type annotation reflect anatomical boundaries of the hippocampal subregions. **(A, B)** UMAP plot **(A)** and spatial transcriptomic maps of hippocampal spots **(B)** from trained and untrained mouse samples. Color reflects clustering of hippocampal spots. **(C, D)** UMAP plot **(C)** and cell-type annotated spatial transcriptomic maps of hippocampal spots **(D)** in samples from the trained and untrained mouse.

Differential gene expression analysis between trained and untrained mice identified total 352 DEGs in all hippocampal spots, with further stratification by subregional cell layers detecting 75 DEGs in the CA1, 72 DEGs in the CA3, and 187 DEGs in the DG (Figure 5 A-E & Supplemental Figure 2). GO enrichment analyses of the DEGs detected between all hippocampal spots identified biological processes related to energy production and protein expression (Figure 5F). DEGs detected in the analysis of the CA1 cell layer were enriched with biological processes related to energy production and synaptic function. In the CA3, DEGs were enriched with genes involved in neurotransmitter release and energy production. And in the DG, DEGs were enriched with genes involved in energy production and translational machinery. Notably, the biological process “regulation of synaptic plasticity” (GO:0048167) was enriched among the DEGs detected between all subregional comparisons between trained and untrained animals. This spatial transcriptomics finding exhibit strong concordance with enrichment patterns from bulk RNA sequencing data, reinforcing a subregion-specific model of memory-associated transcriptional activation.

**Figure 5.**
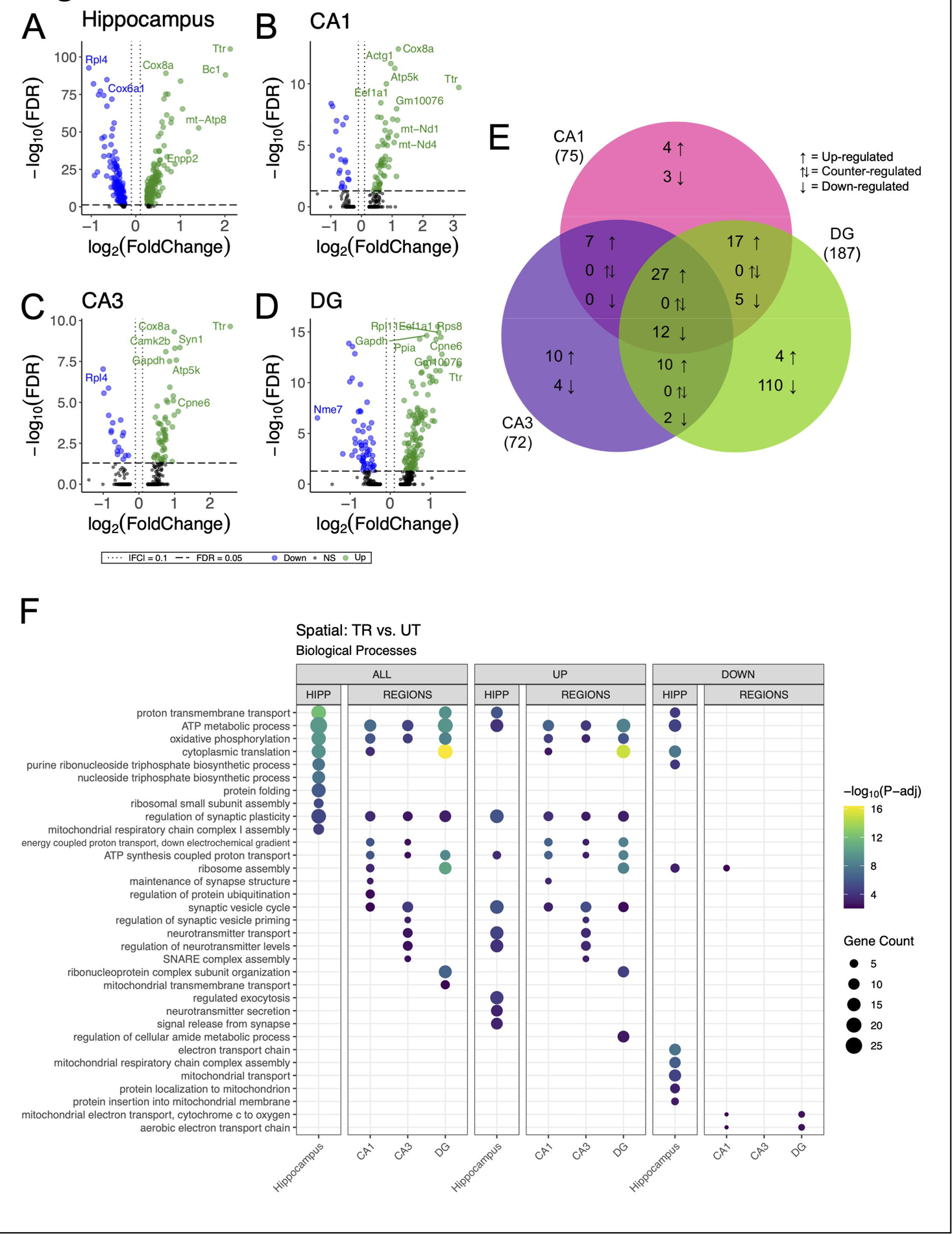
DEGs detected between trained and untrained animals within hippocampal subregions using spatial transcriptomics are enriched with genes involved in similar biological processes to those seen in bulk RNA-seq. **(A-D)** Volcano plot of DEGs between all hippocampal **(A)**, CA1 **(B)**, CA3 **(C)**, and DG **(D)** cell-layer spots from trained and untrained mice. In the analysis of all hippocampal combined, 205 genes were upregulated and 147 were downregulated. 55 genes were upregulated and 20 were downregulated in the CA1 cell layer, 54 genes were upregulated and 18 were downregulated in the CA3 cell layer, and 129 genes were upregulated and 58 were downregulated in the DG cell layer. **(E)** Overlap of regionally detected DEGs stratified by direction of fold change. Overlaps were tested with Fisher’s exact test. See Supplemental Figure 2 for further details. **(F)** Regional enrichment of biological processes detected amongst all DEGs (left) and stratified by up-(middle) and down-regulated (right) DEGs. Dot color reflects the statistical significance (-log_10_(FDR)) of the biological process enrichment. Dot size reflects the number of detected DEGs mapped to the genes involved in a given biological process. See Supplemental Figure 2 for enrichment of Cellular Component and Molecular Function GO terms.

### 3.5 Arc-expressing hippocampal spots exhibit changes in the expression of memory-associated genes and biological processes

Our data suggest that differences in behavioral conditioning contributed to variations in the spatial distribution of synaptic plasticity related gene expression across the hippocampal network. Differences in behavioral conditioning affect the recruitment of *Arc*-expressing hippocampal neurons which form the memory-associated neuronal ensemble (Carrillo-Reid, 2022, Gouty-Colomer et al., 2016, Hsieh et al., 2021, Chia and Otto, 2013). We studied the *Arc*-expressing spatial transcriptomics spots as a proxy for the putative IEG tagged memory-associated neuronal ensemble to assess changes in biological processes following APA memory recall. Out of the 542 spots in the hippocampus from the trained animal, 195 spots had detectable expression of *Arc* mRNA (*Arc+*). The untrained animal, 169 *Arc+* spots out of 568 hippocampal spots (Supplemental Figure 3C). In both samples, spots with the highest level of *Arc* expression concentrated primarily in the CA1 cell layer (Figure 6A, B).

**Figure 6.**
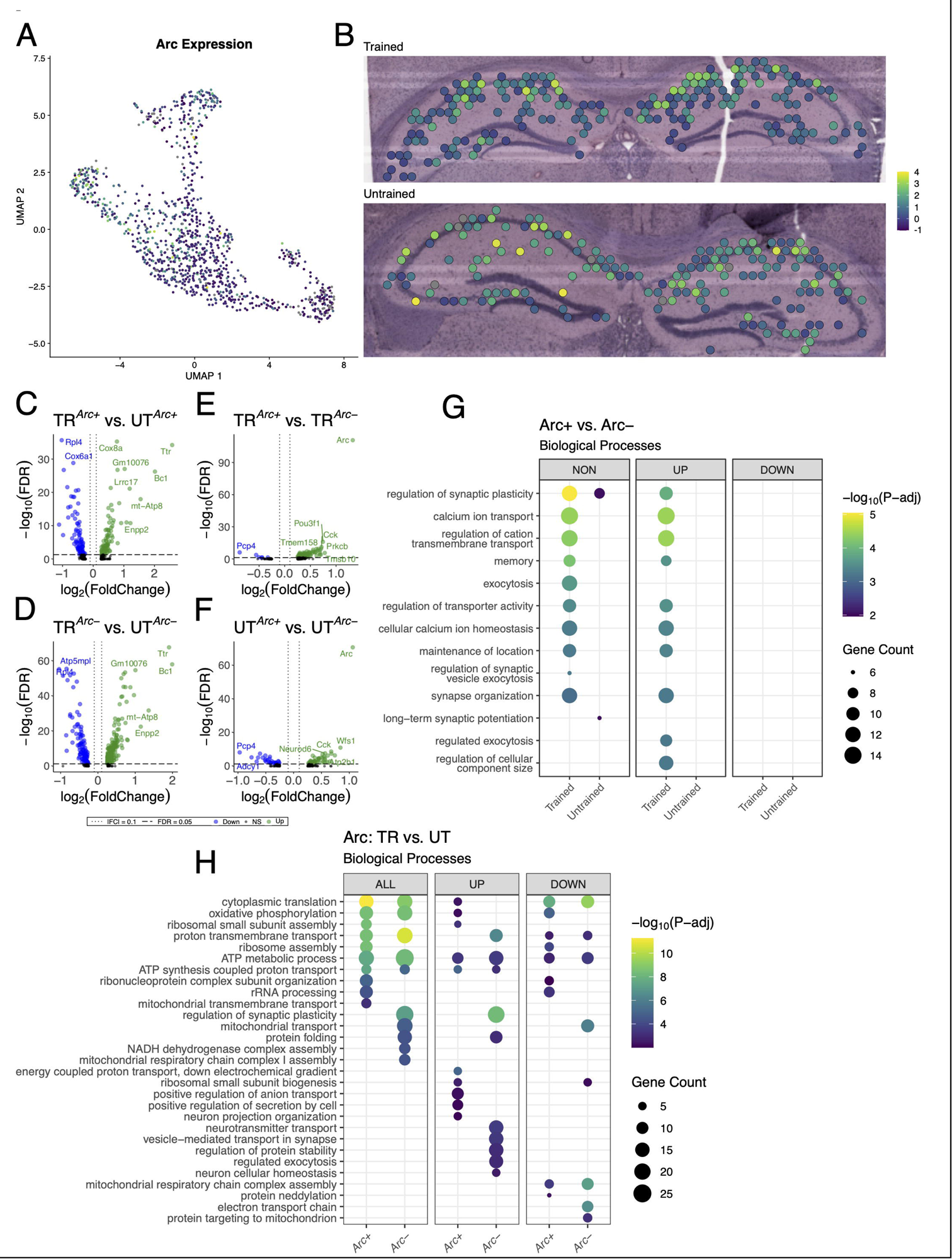
Arc-positive spots in the trained sample are enriched with genes involved in synaptic plasticity. **(A, B)** UMAP plot **(A)** and spatial transcriptomic maps of hippocampal spots **(B)** in the trained and untrained samples colored to reflect normalized expression of *Arc*. **(C, D)** Volcano plots of differential gene expression of *Arc* positive **(C)** and negative **(D)** spots across behavioral conditions. In the analysis of *Arc+* spots across behavioral conditions 85 upregulated and 89 downregulated genes were detected. In the analysis of *Arc-* spots across behavioral conditions 113 upregulated and 240 downregulated genes were detected. **(E, F)** Volcano plots of differential gene expression between *Arc* positive and negative spots within trained **(E)** and untrained **(F)** samples. In the analysis of *Arc* spots within the trained animal 95 upregulated and 6 downregulated genes were detected. In the analysis of *Arc* spots within the untrained animal 47 upregulated and 31 downregulated genes were detected. **(G)** Biological processes overrepresented in the analysis of *Arc*-expressing spots within trained and untrained samples amongst all DEGs (left) and stratified by up-(middle) and down-regulated (right) DEGs. Dot color reflects the statistical significance (-log_10_(FDR)) of the biological process enrichment. Dot size reflects the number of detected DEGs mapped to the genes involved in a given biological process. **(H)** Biological processes overrepresented in the analysis of *Arc*-expressing spots across trained and untrained samples amongst all DEGs (left) and stratified by up-(middle) and down-regulated (right) DEGs.

To assess synaptic plasticity-related transcriptional changes in these *Arc+* spots, we conducted multiple differential expression comparisons: 1) across sets of *Arc+* or *Arc-* spots between the trained and untrained animal (Figure 6C, D), and 2) within the trained or untrained animal between sets of *Arc+* and *Arc-* (Figure 6E, F). The comparison of *Arc+* spots across trained and untrained animals detected 174 significant DEGs, and comparison of *Arc-* spots across trained and untrained animals detected 353 significant DEGs. GO term enrichment analysis of DEGs detected across *Arc*+ spots in the trained and untrained animal revealed enrichment with biological processes related to ribosomal function and energy production (Figure 6H). *Rpl4* and *Rpl5* were differentially expressed in the analysis across *Arc*+ spots in the trained and untrained, and are known for their roles in ribosomal biogenesis (Robledo et al., 2008). Enrichment analyses of DEGs detected in *Arc-* spots across training conditions revealed a similar set biological processes related to energy production as those detected in the analysis of Arc+ spots. Differences between the two comparisons are accentuated when enrichment analyses are performed separately on up- and down-regulated DEGs.

101 DEGs were detected within *Arc*+ and *Arc*-spots in the trained animal, and 78 DEGs were detected within *Arc*+ and *Arc*-spots in the untrained animal. Notably, substantially more biological processes related to synaptic plasticity and neuronal excitability were enriched in the DEGs *Arc* spots of the trained animal as compared to *Arc* spots of the untrained animal. This difference is enhanced when the enrichment analyses are separated by up- and down-regulated DEGs, where only the upregulated DEGs within the *Arc* spots of the trained animal demonstrate enrichment of any biological processes (Figure 6G). *Prkcb* and *Dkk3*, known for their role in synaptic plasticity (Weeber et al., 2000, Martin-Flores et al., 2023), were detected amongst upregulated genes in the comparison of *Arc+* and *Arc*-spots in the trained animal. In summary, enrichment analyses in Arc+ spots across training conditions detected biological processes related to energy metabolism and ribosomal biogenesis, while Arc+ spots within the trained sample revealed biological processes related to synaptic plasticity and transmission.

### 3.6 IEG-expressing hippocampal spots comprise distinct gene expression profiles

Our findings demonstrate that behavioral training induces discrete transcriptional changes in the population of spots marked by *Arc* expression. However, memory traces are likely comprise overlapping yet molecularly distinct neuronal populations tagged by IEGs (Sun et al., 2020, Nambu et al., 2022, Okuno, 2011, Gallo et al., 2018). We sought to examine spots marked by the expression of the IEG most and least correlated with the expression of Arc to evaluate the functional relationship between these subsets of the memory-associated ensemble. Pairwise correlations identified *Egr1* and *c-Jun* as the IEGs most and least correlated with *Arc*, respectively (Supplemental Figure 3 A, B). As such, we performed differential gene expression analyses on *Egr1* and *c-Jun* expressing spots across and within behavioral conditions.

In total, 380 out of 542 spots in the hippocampus of the trained animal and 256 out of 568 spots in the untrained animal were found to express detectable levels of *Egr1* (Supplemental Figure 3G). In both samples, spots with the highest level of *Egr1* expression were located predominantly in the CA1 cell layer, following the spatial distribution of *Arc* expression (Figure 7A, B). Detectable levels of *c-Jun* were seen in 308 out of 542 spots in the hippocampal sample from the trained animal and 175 out of 568 spots in the untrained animal (Supplemental Figure 3G). In both samples, spots with the highest level of *c-Jun* expression were located predominantly in the DG granule cell layer (Figure 7C, D).

**Figure 7.**
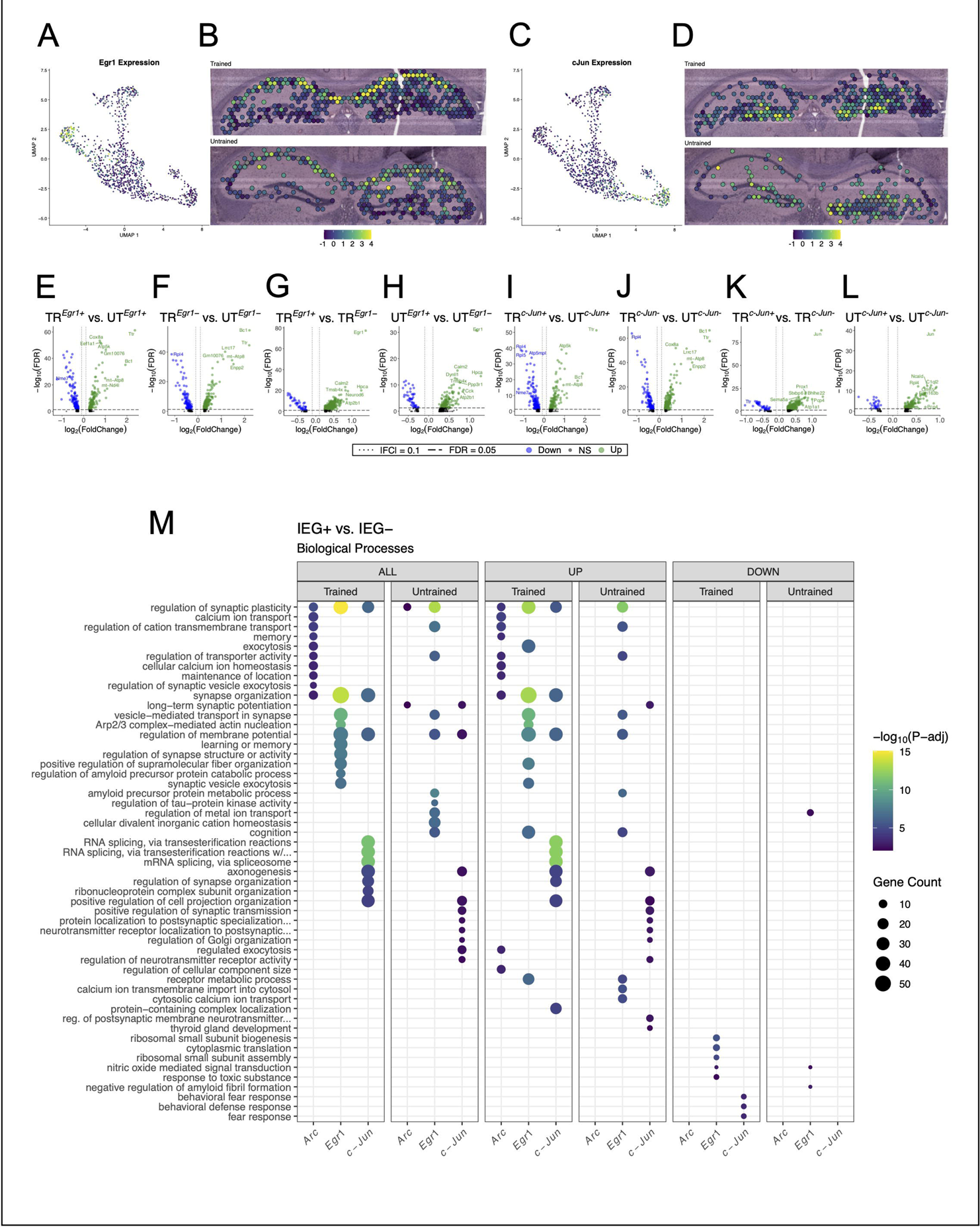
Spatial transcriptomics reveals distinct collections of biological processes enriched among *Arc*, *Egr1*, and *c-Jun*-expressing spots. **(A, B)** UMAP plot **(A)** and spatial transcriptomic maps **(B)** of hippocampal spots in the trained and untrained sample colored to reflect the normalized expression of *Egr1*. **(C, D)** UMAP plot **(C)** and spatial transcriptomic maps **(D)** of hippocampal spots in the trained and untrained samples colored to reflect normalized expression of *c-Jun*. **(E-H)** Volcano plots of differential gene expression of *Egr1* positive **(E)** and negative **(F)** spots across behavioral conditions and within trained **(G)** and untrained **(H)** samples. In the analysis of *Egr1+* spots across behavioral conditions 135 upregulated and 160 downregulated genes were detected. In the analysis of *Egr1-* spots across behavioral conditions 216 upregulated and 111 downregulated genes were detected. In the analysis of *Egr1* spots within the trained animal 509 upregulated and 46 downregulated genes were detected. In the analysis of *Egr1* spots within the untrained animal 150 upregulated and 46 downregulated genes were detected. **(I-L)** Volcano plots of differential gene expression of *c-Jun* positive **(I)** and negative **(J)** spots across behavioral conditions and within trained **(K)** and untrained **(L)** samples. In the analysis of *c-Jun+* spots across behavioral conditions 123 upregulated and 175 downregulated genes were detected. In the analysis of *c-Jun-* spots across behavioral conditions 193 upregulated and 121 downregulated genes were detected. In the analysis of *c-Jun* spots within the trained animal 440 upregulated and 40 downregulated genes were detected. In the analysis of *c-Jun* spots within the untrained animal 138 upregulated and 28 downregulated genes were detected. M) Biological processes overrepresented between the IEG+ and IEG-spots within each sample. Analysis of GO term overrepresentation is stratified by all DEGs (left), up-(middle), and down-regulated (right) DEGs. Dot color reflects statistical significance (-log10(FDR)) of biological process enrichment. Dot size reflects number of detected DEGs mapped to the genes involved in a given biological process.

In the analysis of *Egr1+* or *Egr1-* spots across training conditions, 295 DEGs were detected in the *Egr1+* spots, and 327 DEGs in the *Egr-* spots (Figure 7E, F). In the analysis of *c-Jun+* or *c-Jun-* spots across training conditions, 298 DEGs were detected in the *c-Jun +* spots, and 314 DEGs in the *c-Jun-* spots (Figure 7I, J). GO enrichment analyses of all three IEG expressing spot populations across trained and untrained conditions revealed similar sets of enriched biological processes (Supplemental Figure 4D), namely those involved in ribosomal function and energy production.

In the analysis between *Egr1+* and *Egr-* spots, 555 DEGs were identified within the trained animal, and 196 DEGs within the untrained animal (Figure 7G, H). In the analysis between *c-Jun+* and *c-Jun*-spots, 480 DEGs were identified within the trained animal, and 166 DEGs within the untrained animal (Figure 7K, L).

Due to the high degree of correlation between the expression of *Arc* and *Egr1*, and the low correlation between the expression of *Arc* and *c-Jun*, we expected that *Arc* and *Egr1*-expressing spots would appear more functionally similar to one another than they would to *c-Jun*-expressing spots, within the same sample. Counterintuitively, comparisons of IEG+ and IEG-spots within each sample revealed distinct collections of biological processes overrepresented in the *Arc*, *Egr1*, and *c-Jun* expressing populations. Additionally, in contrast to the within sample analyses of *Arc* spots (Figure 6G), within sample analyses of *Egr1* and *c-Jun* spots were enriched with numerous biological processes in *both* trained and untrained mice. This effect is enhanced when enrichment analyses are stratified by up- and down-regulation. Biological processes enriched among DEGs in the *Egr1* spots within the trained and untrained mice were both involved in synaptic organization and plasticity (Figure 7M). Biological processes involved in RNA splicing, synaptic plasticity, and axonal growth were overrepresented in *c-Jun* spots in both trained and untrained mice. Together, these data suggest that IEG-expression following memory recall marks subsets of neurons which are enriched with distinct sets of biological processes.

### 3.7 RT-qPCR validation of regionally detected DEGs

We performed targeted gene expression analyses using RT-qPCR on DG, CA3, and CA1 subregion samples collected from similarly trained animals (Supplemental Figure 4A). Based on region specific DEG identification in the bulk RNA sequencing study, we selected *Kit* (DG), *Sst* (CA3), *Cdh24* (CA1), and *Slc6a7* (overlap of all regions) as representative targets, with *Gapdh* as an internal control. In trained mice, significant upregulation of *Kit* and *Sst* was detected in the DG (Supplemental Figure 4D). Analyses in the CA3 subregion were unable to detect significant differences in gene expression (Supplemental Figure 4C). In the CA1 subregion, increased expression of *Cdh24* was noted, but it did not reach statistical significance (Supplemental Figure 4B). *Slc6a7* was unchanged in all subregions. In summary, RT-qPCR provides a partial validation of our RNA sequencing findings, confirming the upregulation of top DEGs *Kit* and *Sst* selectively in the DG of trained mice.

## 4 DISCUSSION

In this study, we used a combination of bulk RNA sequencing and spatial transcriptomics to identify changes in gene expression within each major subregion of the dorsal hippocampus and within IEG-expressing spots following memory recall. A recent study utilizing spatial transcriptomics found distinct transcriptomic signatures in the hippocampal subregions following training in a spatial object recognition task (Vanrobaeys et al., 2023). This finding illustrates the power of using novel transcriptomic analyses to explore the spatial distribution of gene expression profiles when investigating memory-associated neuronal networks.

### 4.1 Regionalization of molecular mechanisms following memory recall

Different brain regions are recruited depending on the type of memory training an animal is exposed to (Rolls, 2000, Tonegawa et al., 2018). We found evidence supporting the regionalization of gene expression resulting in overrepresentation of GO terms across the dorsal hippocampus following spatial memory recall. A finding consistent with prior reports demonstrating enrichment of biological processes related to transcription and synaptic differentiation in the DG following APA recall (Harris et al., 2020). In our study, comparative bulk RNA sequencing analyses between trained and untrained animals revealed abundant overlap of detected DEGs and enriched biological processes between the CA1 and CA3 subregions. However, the transcriptional landscape observed in the DG exhibited divergence from both CA subregions, suggesting recruitment of specialized molecular pathways in a subfield specific manner following memory recall.

Cells in the CA1 and CA3 subregions share similar morphologies and ontology as compared to the dentate gyrus (Stanfield and Cowan, 1979, Khalaf-Nazzal and Francis, 2013, Cembrowski and Spruston, 2019), which could account for our regionalized findings in the bulk RNA sequencing analysis. During memory recall, the functional coupling of the CA3 and CA1 subregions (Montgomery and Buzsaki, 2007, Carr et al., 2012) is also reflected by the gene expression changes observed in this analysis. Our findings support the role of neurogenesis and protein synthesis in the DG (Alam et al., 2018, Kitamura et al., 2009, Kitamura and Inokuchi, 2014, Deng et al., 2010), as well as the memory-associated increases in efficacy in CA3-CA1 synaptic connections (Aksoy-Aksel and Manahan-Vaughan, 2013, Pavlowsky and Alarcon, 2012). Furthermore, our data also implicate enhanced energetic requirements in the DG during periods of high cognitive demand that enable accurate spatial navigation and memory recall (Mendez-Lopez et al., 2010, Christie and Schrater, 2015), evidenced by the enrichment of ATP metabolic pathways.

### 4.2 Technical considerations of using spatial transcriptomics

We employed an integrative approach combining bulk RNA sequencing and spatial transcriptomics to determine the transcriptomic profile elicited in the hippocampus following the recall of a consolidated spatial memory. Both methodologies provided unique utility and insight into the transcriptomic changes in the brain following memory and have proven to be highly compatible (Li and Wang, 2021, Vanrobaeys et al., 2023). Our bulk RNA sequencing uncovered robust hippocampal subregion-specific distinctions in biological process enrichment between trained and untrained conditions. Meanwhile, spatial transcriptomic analyses revealed anatomical stratification of DEGs but convergence on overlapping sets of enriched biological processes across the CA1, CA3, and DG cell layers. These two distinct outcomes potentially arise from key chemical and technical variances between the two methodologies. First, the 55µm capture spots of the 10x Visium spatial transcriptomics platform permits a more granular analysis of gene expression differences between specifically annotated populations of cells, as opposed to the entirety of a bulk section (Stahl et al., 2016). Second, relative to bulk sequencing, 10x’s Visium spatial transcriptomics requires more tissue processing before RNA extraction and has lower RNA detection sensitivity (Asp et al., 2020), which might skew the genes detected between the two studies. Nevertheless, our dual omics approach leverages the combined strength and resolution of these emerging technologies to simultaneously explore the gene expression changes following spatial memory recall within individual cell populations across a spatially distributed neuronal network.

### 4.3 *Arc*-expressing spots are enriched with genes involved in synaptic plasticity

Using spatial transcriptomics, we examined the spatial distribution of IEG expression in the dorsal hippocampus to infer the location of neurons involved in memory formation. We hypothesized that spatially restricted subsets of IEG-expressing memory-associated neurons would be detected amongst these spots and enriched with genes related to synaptic plasticity. To test this, mice were sacrificed following retention test during the temporal window comprising the peak expression of IEGs as well as the initial upregulation phase of late response genes (Yap and Greenberg, 2018, Sheng and Greenberg, 1990). This approach enabled concurrent transcriptional profiling of both the rapid and delayed genomic responses linked to memory formation. The spatial patterning of IEG induction and overlap with plasticity-related genes could reveal the locations of neurons recruited to the memory trace with the dorsal hippocampus.

Comparisons of *Arc*+ spots across the trained and untrained animal revealed enrichment of biological processes related to energy metabolism and ribosomal function. Interestingly, comparisons of *Arc*+ versus *Arc*-spots within the trained animal were substantially different from those in the untrained animal. Specifically, *Arc*+ spots in the trained animal displayed upregulation of biological processes related to synaptic plasticity, several of which were also enriched in the bulk analyses. These data imply that the significant functional modulation occurring selectively the *Arc*-expressing neuronal population following learning may largely drive the transcriptional differences observed between training conditions in the analysis of bulk hippocampal subregions.

It is not yet known if the functional changes that distinguish cells in a memory-associated neuronal ensemble from surrounding cells are the same functional changes that distinguish these cells across training conditions. Our data suggests that there are two defining features of the *Arc+* spots: the upregulation of synaptic plasticity mechanisms relative to *Arc*-spots, and the upregulation of energy production and ribosomal function in the trained versus the untrained animal. In support of these findings, a recent study identified synaptic plasticity as a defining feature of a neuron in a memory-associated neuronal ensemble (Jeong et al., 2021). Additionally, increases in energy production and ribosomal biogenesis have been found during periods of high cognitive demand (Magistretti and Allaman, 2015), (Hernández et al., 2015), which corresponds the trained avoidance behavior of the APA paradigm (Cimadevilla et al., 2000, Wesierska et al., 2005).

### 4.4 *Egr1-* and *c-Jun*-expressing spots exhibit distinct transcriptomic profiles

Extensively studied for its role in memory storage, the Arc-tagged neuronal ensemble represents only a fraction of neurons activated during a memory event (Lacagnina et al., 2019, Sun et al., 2020, Sweis et al., 2021). Distinct subgroups of the neurons activated during a memory event are marked by several Immediate Early Genes (IEGs), which are associated with different patterns of intense neuronal activity (Okuno, 2011, Sun and Lin, 2016, Sheng et al., 1993, Guzowski et al., 2001, Tonegawa et al., 2018). This IEG expressing neuronal ensembles could encode multiple memory traces for a specific behavioral experience (Kitamura et al., 2017, Tonegawa et al., 2018, Tonegawa et al., 2015, Terranova et al., 2023). Using spatial transcriptomics, we assessed the gene expression profiles of spots expressing the IEGs *Egr1* and *c-Jun*, which were the IEGs most and least correlated with *Arc* expression, respectively.

While partially overlapping, *Arc+, Egr1+,* and *c-Jun+* spots displayed enrichment for distinct sets of biological processes within both trained and untrained animals. In contrast to the within sample comparisons of *Arc* spots, numerous biological processes were overrepresented within the trained and untrained animal for both *Egr1* and *c-Jun* spots. Specifically, *Egr1* spots displayed enrichment of biological processes related to synaptic organization and synaptic plasticity, whereas *c-Jun* spots displayed enrichment of biological processes related to RNA splicing, synaptic plasticity, and axonal projection. Additionally, the spatial distribution of *Arc* and *Egr1* shows high levels of expression in the CA1 cell layer of trained and untrained samples, while the expression of *c-Jun* is high in the DG.

Differing IEG induction likely relates to unique patterns of neuronal activation (Lyons and West, 2011, Madabhushi and Kim, 2018, Tyssowski et al., 2018, Flavell and Greenberg, 2008) and downstream circuit engagement (Roy et al., 2017), highlighting the functional and spatial diversity amongst recruited cells within memory associated neuronal ensembles. Of note, *Egr1* and *c-Jun* expression has been reported in response to fear memory and acute stress, respectively (Cheval et al., 2012, Maddox et al., 2011, Sherrin et al., 2010). The differences observed amongst the *Egr1* and *c-Jun* expressing spots could be related to emotional, temporal, and contextual components inherent to both long term memory experiences of trained and untrained animals in the APA apparatus.

Unlike *within* sample analyses, cross condition comparison revealed consistent enrichment for biological processes related to energy production across the IEG+ and IEG-spots, fitting with the heightened metabolic activity previously observed during periods of increased cognitive demand (Christie and Schrater, 2015, Magistretti and Allaman, 2015).

### 4.5 Investigating memory across multiple scales in the brain

In summary, we performed an integrated investigation of hippocampal transcriptional dynamics following spatial memory recall using bulk RNA sequencing and spatial transcriptomics. Regionally restricted expression patterns were observed with the CA1 and CA3 subregion exhibiting enrichment for synaptic plasticity and transmission pathways, while the DG was more prominently categorized by enrichment for protein synthesis and energy metabolism pathways. We identified a specialized signature of *Arc* expressing spots in trained mice categorized by upregulation of genes involved synaptic plasticity and transmission. Additionally, functionally distinct IEG expressing populations were revealed with *Arc, Egr1,* and *c-Jun* expressing spots exhibiting differential pathway enrichment and anatomical distribution.

The hippocampal transcriptional landscape captured in this study of APA memory recall represents a single transitionary state over the life of a memory. Networks in the hippocampus undergo complex spatio-temporal tuning in flow of information depending on memory phase, valence, and cognitive load (Terranova et al., 2022, Roy et al., 2017, Marks et al., 2022). As a memory evolves through systems consolidation, the neural ensembles, cellular properties, and molecular profiles supporting the memory trace are thought to transform correspondingly (Tonegawa et al., 2018, Alberini and Kandel, 2014). However, capturing this gradual reshaping of memory representations across hippocampal subregions has remained challenging. Emerging spatial molecular profiling techniques offer unmatched resolution for mapping distributed neuronal populations while preserving native tissue context. By applying transcriptomic methods to concurrently inspect spatial loci and neuronal populations, the topological features of the molecular signatures supporting memory could be elucidated.

## Supporting information

Supplemental Table 1. Differential Gene Expression Tables

Supplemental Table 2. GO Enrichment Analysis Tables

## ACKNOWLEDGEMENTS

Supported by Senior Vice President of Research Seed Grant (State University of New York, Downstate Health Sciences University) and partial support from NINDS grant R21NS091830.

## REFERENCES

10X Genomics 2023. Visium Spatial Protocols – Tissue Preparation Guide.

Aksoy-Aksel, A. & Manahan-Vaughan, D. 2013. The temporoammonic input to the hippocampal CA1 region displays distinctly different synaptic plasticity compared to the Schaffer collateral input in vivo: significance for synaptic information processing. Front Synaptic Neurosci, 5, 5.

Alam, M. J., Kitamura, T., Saitoh, Y., Ohkawa, N., Kondo, T. & Inokuchi, K. 2018. Adult Neurogenesis Conserves Hippocampal Memory Capacity. J Neurosci, 38, 6854–6863.

Alberini, C. M. & Kandel, E. R. 2014. The regulation of transcription in memory consolidation. Cold Spring Harb Perspect Biol, 7, a021741.

Amaral, D. G. & Witter, M. P. 1989. The three-dimensional organization of the hippocampal formation: a review of anatomical data. Neuroscience, 31, 571–91.

Annamneedi, A., Del Angel, M., Gundelfinger, E. D., Stork, O. & Caliskan, G. 2021. The Presynaptic Scaffold Protein Bassoon in Forebrain Excitatory Neurons Mediates Hippocampal Circuit Maturation: Potential Involvement of Trkb Signalling. Int J Mol Sci, 22.

Asok, A., Leroy, F., Rayman, J. B. & Kandel, E. R. 2019. Molecular Mechanisms of the Memory Trace. Trends Neurosci, 42, 14–22.

Asp, M., Bergenstrahle, J. & Lundeberg, J. 2020. Spatially Resolved Transcriptomes-Next Generation Tools for Tissue Exploration. Bioessays, 42, e1900221.

Barbieri, R., Contestabile, A., Ciardo, M. G., Forte, N., Marte, A., Baldelli, P., Benfenati, F. & Onofri, F. 2018. Synapsin I and Synapsin II regulate neurogenesis in the dentate gyrus of adult mice. Oncotarget, 9, 18760–18774.

Basu, J. & Siegelbaum, S. A. 2015. The Corticohippocampal Circuit, Synaptic Plasticity, and Memory. Cold Spring Harb Perspect Biol, 7.

Butler, A., Hoffman, P., Smibert, P., Papalexi, E. & Satija, R. 2018. Integrating single-cell transcriptomic data across different conditions, technologies, and species. Nat Biotechnol, 36, 411–420.

Cai, H., Chen, H., Yi, T., Daimon, C. M., Boyle, J. P., Peers, C., Maudsley, S. & Martin, B. 2013. VennPlex–A Novel Venn Diagram Program for Comparing and Visualizing Datasets with Differentially Regulated Datapoints. PLOS ONE, 8, e53388.

Carr, M. F., Karlsson, M. P. & Frank, L. M. 2012. Transient slow gamma synchrony underlies hippocampal memory replay. Neuron, 75, 700–13.

Carrillo-Reid, L. 2022. Neuronal ensembles in memory processes. Semin Cell Dev Biol, 125, 136–143.

Cembrowski, M. S. & Spruston, N. 2019. Heterogeneity within classical cell types is the rule: lessons from hippocampal pyramidal neurons. Nat Rev Neurosci, 20, 193–204.

Chen, B. K., Murawski, N. J., Cincotta, C., Mckissick, O., Finkelstein, A., Hamidi, A. B., Merfeld, E., Doucette, E., Grella, S. L., Shpokayte, M., Zaki, Y., Fortin, A. & Ramirez, S. 2019. Artificially Enhancing and Suppressing Hippocampus-Mediated Memories. Curr Biol, 29, 1885–1894 e4.

Cheval, H., Chagneau, C., Levasseur, G., Veyrac, A., Faucon-Biguet, N., Laroche, S. & Davis, S. 2012. Distinctive features of Egr transcription factor regulation and DNA binding activity in CA1 of the hippocampus in synaptic plasticity and consolidation and reconsolidation of fear memory. Hippocampus, 22, 631–42.

Chevaleyre, V. & Siegelbaum, S. A. 2010. Strong CA2 pyramidal neuron synapses define a powerful disynaptic cortico-hippocampal loop. Neuron, 66, 560–572.

Chia, C. & Otto, T. 2013. Hippocampal Arc (Arg3.1) expression is induced by memory recall and required for memory reconsolidation in trace fear conditioning. Neurobiol Learn Mem, 106, 48–55.

Cho, J. H., Huang, B. S. & Gray, J. M. 2016. RNA sequencing from neural ensembles activated during fear conditioning in the mouse temporal association cortex. Sci Rep, 6, 31753.

Christie, S. T. & Schrater, P. 2015. Cognitive cost as dynamic allocation of energetic resources. Front Neurosci, 9, 289.

Cimadevilla, J. M., Kaminsky, Y., Fenton, A. & Bures, J. 2000. Passive and active place avoidance as a tool of spatial memory research in rats. J Neurosci Methods, 102, 155–64.

Collins, B. E., Greer, C. B., Coleman, B. C. & Sweatt, J. D. 2019. Histone H3 lysine K4 methylation and its role in learning and memory. Epigenetics & Chromatin, 12, 7.

Deng, W., Aimone, J. B. & Gage, F. H. 2010. New neurons and new memories: how does adult hippocampal neurogenesis affect learning and memory? Nat Rev Neurosci, 11, 339–50.

Denny, C. A., Kheirbek, M. A., Alba, E. L., Tanaka, K. F., Brachman, R. A., Laughman, K. B., Tomm, N. K., Turi, G. F., Losonczy, A. & Hen, R. 2014. Hippocampal memory traces are differentially modulated by experience, time, and adult neurogenesis. Neuron, 83, 189–201.

Erwin, S. R., Sun, W., Copeland, M., Lindo, S., Spruston, N. & Cembrowski, M. S. 2020. A Sparse, Spatially Biased Subtype of Mature Granule Cell Dominates Recruitment in Hippocampal-Associated Behaviors. Cell Rep, 31, 107551.

Ferreira, E., Shaw, D. M. & Oddo, S. 2016. Identification of learning-induced changes in protein networks in the hippocampi of a mouse model of Alzheimer’s disease. Transl Psychiatry, 6, e849.

Flavell, S. W. & Greenberg, M. E. 2008. Signaling mechanisms linking neuronal activity to gene expression and plasticity of the nervous system. Annu Rev Neurosci, 31, 563–90.

Gallo, F. T., Katche, C., Morici, J. F., Medina, J. H. & Weisstaub, N. V. 2018. Immediate Early Genes, Memory and Psychiatric Disorders: Focus on c-Fos, Egr1 and Arc. Front Behav Neurosci, 12, 79.

Genomics, X. 2019. Visium Spatial Gene Expression Reagent Kits - Tissue Optimization.

Genomics, X. 2021. Visium Spatial Gene Expression Imaging Guidelines.

Genomics, X. 2023. Visium Spatial Gene Expression Reagent Kits User Guide.

Gouty-Colomer, L. A., Hosseini, B., Marcelo, I. M., Schreiber, J., Slump, D. E., Yamaguchi, S., Houweling, A. R., Jaarsma, D., Elgersma, Y. & Kushner, S. A. 2016. Arc expression identifies the lateral amygdala fear memory trace. Mol Psychiatry, 21, 364–75.

Gu, Z., Eils, R. & Schlesner, M. 2016. Complex heatmaps reveal patterns and correlations in multidimensional genomic data. Bioinformatics, 32, 2847–2849.

Guan, J. S., Jiang, J., Xie, H. & Liu, K. Y. 2016. How Does the Sparse Memory “Engram” Neurons Encode the Memory of a Spatial-Temporal Event? Front Neural Circuits, 10, 61.

Guzowski, J. F., Mcnaughton, B. L., Barnes, C. A. & Worley, P. F. 1999. Environment-specific expression of the immediate-early gene Arc in hippocampal neuronal ensembles. Nat Neurosci, 2, 1120–4.

Guzowski, J. F., Setlow, B., Wagner, E. K. & Mcgaugh, J. L. 2001. Experience-dependent gene expression in the rat hippocampus after spatial learning: a comparison of the immediate-early genes Arc, c-fos, and zif268. J Neurosci, 21, 5089–98.

Hao, Y., Stuart, T., Kowalski, M. H., Choudhary, S., Hoffman, P., Hartman, A., Srivastava, A., Molla, G., Madad, S., Fernandez-Granda, C. & Satija, R. 2023. Dictionary learning for integrative, multimodal and scalable single-cell analysis. Nature Biotechnology.

Harris, R. M., Kao, H.-Y., Alarcón, J. M., Fenton, A. A. & Hofmann, H. A. 2020. Transcriptome analysis of hippocampal subfields identifies gene expression profiles associated with long-term active place avoidance memory. bioRxiv, 2020.02.05.935759.

Hernández, A. I., Alarcon, J. M. & Allen, K. D. 2015. New ribosomes for new memories? Communicative & Integrative Biology, 8, e1017163.

Hsieh, C., Tsokas, P., Grau-Perales, A., Lesburgueres, E., Bukai, J., Khanna, K., Chorny, J., Chung, A., Jou, C., Burghardt, N. S., Denny, C. A., Flores-Obando, R. E., Hartley, B. R., Rodriguez Valencia, L. M., Hernandez, A. I., Bergold, P. J., Cottrell, J. E., Alarcon, J. M., Fenton, A. A. & Sacktor, T. C. 2021. Persistent increases of PKMzeta in memory-activated neurons trace LTP maintenance during spatial long-term memory storage. Eur J Neurosci, 54, 6795–6814.

Hunsaker, M. R. & Kesner, R. P. 2013. The operation of pattern separation and pattern completion processes associated with different attributes or domains of memory. Neurosci Biobehav Rev, 37, 36–58.

Irfan, M., Gopaul, K. R., Miry, O., Hökfelt, T., Stanton, P. K. & Bark, C. 2019. Snap-25 isoforms differentially regulate synaptic transmission and long-term synaptic plasticity at central synapses. Scientific Reports, 9, 6403.

Jeong, Y., Cho, H. Y., Kim, M., Oh, J. P., Kang, M. S., Yoo, M., Lee, H. S. & Han, J. H. 2021. Synaptic plasticity-dependent competition rule influences memory formation. Nat Commun, 12, 3915.

Josselyn, S. A. & Tonegawa, S. 2020. Memory engrams: Recalling the past and imagining the future. Science, 367.

Katafuchi, T., Li, A. J., Hirota, S., Kitamura, Y. & Hori, T. 2000. Impairment of spatial learning and hippocampal synaptic potentiation in c-kit mutant rats. Learn Mem, 7, 383–92.

Kesner, R. P., Lee, I. & Gilbert, P. 2004. A behavioral assessment of hippocampal function based on a subregional analysis. Rev Neurosci, 15, 333–51.

Khalaf-Nazzal, R. & Francis, F. 2013. Hippocampal development - old and new findings. Neuroscience, 248, 225–42.

Kitamura, T. & Inokuchi, K. 2014. Role of adult neurogenesis in hippocampal-cortical memory consolidation. Mol Brain, 7, 13.

Kitamura, T., Ogawa, S. K., Roy, D. S., Okuyama, T., Morrissey, M. D., Smith, L. M., Redondo, R. L. & Tonegawa, S. 2017. Engrams and circuits crucial for systems consolidation of a memory. Science, 356, 73–78.

Kitamura, T., Saitoh, Y., Takashima, N., Murayama, A., Niibori, Y., Ageta, H., Sekiguchi, M., Sugiyama, H. & Inokuchi, K. 2009. Adult neurogenesis modulates the hippocampus-dependent period of associative fear memory. Cell, 139, 814–27.

Kouser, M., Speed, H. E., Dewey, C. M., Reimers, J. M., Widman, A. J., Gupta, N., Liu, S., Jaramillo, T. C., Bangash, M., Xiao, B., Worley, P. F. & Powell, C. M. 2013. Loss of predominant Shank3 isoforms results in hippocampus-dependent impairments in behavior and synaptic transmission. J Neurosci, 33, 18448–68.

Lacagnina, A. F., Brockway, E. T., Crovetti, C. R., Shue, F., Mccarty, M. J., Sattler, K. P., Lim, S. C., Santos, S. L., Denny, C. A. & Drew, M. R. 2019. Distinct hippocampal engrams control extinction and relapse of fear memory. Nat Neurosci, 22, 753–761.

Lee, B. H., Shim, J. Y., Moon, H. C., Kim, D. W., Kim, J., Yook, J. S., Kim, J. & Park, H. Y. 2022. Real-time visualization of mRNA synthesis during memory formation in live mice. Proc Natl Acad Sci U S A, 119, e2117076119.

Leone, L., Colussi, C., Gironi, K., Longo, V., Fusco, S., Li Puma, D. D., D’ascenzo, M. & Grassi, C. 2019. Altered Nup153 Expression Impairs the Function of Cultured Hippocampal Neural Stem Cells Isolated from a Mouse Model of Alzheimer’s Disease. Mol Neurobiol, 56, 5934–5949.

Li, X. & Wang, C. Y. 2021. From bulk, single-cell to spatial RNA sequencing. Int J Oral Sci, 13, 36.

Lisman, J. E. 1999. Relating hippocampal circuitry to function: recall of memory sequences by reciprocal dentate-CA3 interactions. Neuron, 22, 233–42.

Lyons, M. R. & West, A. E. 2011. Mechanisms of specificity in neuronal activity-regulated gene transcription. Prog Neurobiol, 94, 259–95.

Madabhushi, R. & Kim, T. K. 2018. Emerging themes in neuronal activity-dependent gene expression. Mol Cell Neurosci, 87, 27–34.

Maddox, S. A., Monsey, M. S. & Schafe, G. E. 2011. Early growth response gene 1 (Egr-1) is required for new and reactivated fear memories in the lateral amygdala. Learn Mem, 18, 24–38.

Magee, J. C. & Grienberger, C. 2020. Synaptic Plasticity Forms and Functions. Annual Review of Neuroscience, 43, 95–117.

Magistretti, P. J. & Allaman, I. 2015. A cellular perspective on brain energy metabolism and functional imaging. Neuron, 86, 883–901.

Malenka, R. C. & Bear, M. F. 2004. LTP and LTD: an embarrassment of riches. Neuron, 44, 5–21.

Mao, W., Watanabe, T., Cho, S., Frost, J. L., Truong, T., Zhao, X. & Futai, K. 2015. Shank1 regulates excitatory synaptic transmission in mouse hippocampal parvalbumin-expressing inhibitory interneurons. Eur J Neurosci, 41, 1025–35.

Marco, A., Meharena, H. S., Dileep, V., Raju, R. M., Davila-Velderrain, J., Zhang, A. L., Adaikkan, C., Young, J. Z., Gao, F., Kellis, M. & Tsai, L. H. 2020. Mapping the epigenomic and transcriptomic interplay during memory formation and recall in the hippocampal engram ensemble. Nat Neurosci, 23, 1606–1617.

Marks, W. D., Yokose, J., Kitamura, T. & Ogawa, S. K. 2022. Neuronal Ensembles Organize Activity to Generate Contextual Memory. Front Behav Neurosci, 16, 805132.

Martin-Flores, N., Podpolny, M., Mcleod, F., Workman, I., Crawford, K., Ivanov, D., Leonenko, G., Escott-Price, V. & Salinas, P. C. 2023. Downregulation of Dickkopf-3, a Wnt antagonist elevated in Alzheimer’s disease, restores synapse integrity and memory in a disease mouse model. bioRxiv, 2022.06.16.495307.

Mayford, M. 2014. The search for a hippocampal engram. Philos Trans R Soc Lond B Biol Sci, 369, 20130161.

Mayford, M., Siegelbaum, S. A. & Kandel, E. R. 2012. Synapses and memory storage. Cold Spring Harb Perspect Biol, 4.

Meenakshi, P., Kumar, S. & Balaji, J. 2021. In vivo imaging of immediate early gene expression dynamics segregates neuronal ensemble of memories of dual events. Mol Brain, 14, 102.

Mendez-Lopez, M., Mendez, M., Begega, A. & Arias, J. L. 2010. Spatial short-term memory in rats: effects of learning trials on metabolic activity of limbic structures. Neurosci Lett, 483, 32–5.

Middleton, S. J. & Mchugh, T. J. 2020. CA2: A Highly Connected Intrahippocampal Relay. Annu Rev Neurosci, 43, 55–72.

Molitor, R. J., Sherrill, K. R., Morton, N. W., Miller, A. A. & Preston, A. R. 2021. Memory Reactivation during Learning Simultaneously Promotes Dentate Gyrus/CA_2,3_ Pattern Differentiation and CA_1_ Memory Integration. The Journal of Neuroscience, 41, 726–738.

Montgomery, S. M. & Buzsaki, G. 2007. Gamma oscillations dynamically couple hippocampal CA3 and CA1 regions during memory task performance. Proc Natl Acad Sci U S A, 104, 14495–500.

Nambu, M. F., Lin, Y. J., Reuschenbach, J. & Tanaka, K. Z. 2022. What does engram encode?: Heterogeneous memory engrams for different aspects of experience. Curr Opin Neurobiol, 75, 102568.

Okuno, H. 2011. Regulation and function of immediate-early genes in the brain: beyond neuronal activity markers. Neurosci Res, 69, 175–86.

Pavlowsky, A. & Alarcon, J. M. 2012. Interaction between long-term potentiation and depression in CA1 synapses: temporal constrains, functional compartmentalization and protein synthesis. PLoS One, 7, e29865.

Paxinos, G. & Franklin, K. B. J. 2019. Paxinos and Franklin’s the Mouse Brain in Stereotaxic Coordinates, Elsevier Science.

Ponchel, F., Toomes, C., Bransfield, K., Leong, F. T., Douglas, S. H., Field, S. L., Bell, S. M., Combaret, V., Puisieux, A., Mighell, A. J., Robinson, P. A., Inglehearn, C. F., Isaacs, J. D. & Markham, A. F. 2003. Real-time PCR based on SYBR-Green I fluorescence: an alternative to the TaqMan assay for a relative quantification of gene rearrangements, gene amplifications and micro gene deletions. BMC Biotechnol, 3, 18.

Poo, M. M., Pignatelli, M., Ryan, T. J., Tonegawa, S., Bonhoeffer, T., Martin, K. C., Rudenko, A., Tsai, L. H., Tsien, R. W., Fishell, G., Mullins, C., Goncalves, J. T., Shtrahman, M., Johnston, S. T., Gage, F. H., Dan, Y., Long, J., Buzsaki, G. & Stevens, C. 2016. What is memory? The present state of the engram. BMC Biol, 14, 40.

Ramirez-Amaya, V., Vazdarjanova, A., Mikhael, D., Rosi, S., Worley, P. F. & Barnes, C. A. 2005. Spatial exploration-induced Arc mrna and protein expression: evidence for selective, network-specific reactivation. J Neurosci, 25, 1761–8.

Rao-Ruiz, P., Couey, J. J., Marcelo, I. M., Bouwkamp, C. G., Slump, D. E., Matos, M. R., Van Der Loo, R. J., Martins, G. J., Van Den Hout, M., Van, I. W. F., Costa, R. M., Van Den Oever, M. C. & Kushner, S. A. 2019. Engram-specific transcriptome profiling of contextual memory consolidation. Nat Commun, 10, 2232.

Reshetnikov, V. V., Kisaretova, P. E., Ershov, N. I., Shulyupova, A. S., Oshchepkov, D. Y., Klimova, N. V., Ivanchihina, A. V., Merkulova, T. I. & Bondar, N. P. 2020. Genes associated with cognitive performance in the Morris water maze: an Rna-seq study. Sci Rep, 10, 22078.

Robledo, S., Idol, R. A., Crimmins, D. L., Ladenson, J. H., Mason, P. J. & Bessler, M. 2008. The role of human ribosomal proteins in the maturation of rRNA and ribosome production. RNA, 14, 1918–29.

Rolls, E. T. 2000. Memory systems in the brain. Annu Rev Psychol, 51, 599–630.

Roy, D. S., Kitamura, T., Okuyama, T., Ogawa, S. K., Sun, C., Obata, Y., Yoshiki, A. & Tonegawa, S. 2017. Distinct Neural Circuits for the Formation and Retrieval of Episodic Memories. Cell, 170, 1000–1012 e19.

Ryan, T. J., Roy, D. S., Pignatelli, M., Arons, A. & Tonegawa, S. 2015. Memory. Engram cells retain memory under retrograde amnesia. Science, 348, 1007–13.

Sanes, J. R. & Lichtman, J. W. 1999. Can molecules explain long-term potentiation? Nat Neurosci, 2, 597–604.

Sauvage, M., Kitsukawa, T. & Atucha, E. 2019. Single-cell memory trace imaging with immediate-early genes. J Neurosci Methods, 326, 108368.

Senzai, Y. 2019. Function of local circuits in the hippocampal dentate gyrus-CA3 system. Neurosci Res, 140, 43–52.

Shah, S., Lubeck, E., Zhou, W. & Cai, L. 2016. In Situ Transcription Profiling of Single Cells Reveals Spatial Organization of Cells in the Mouse Hippocampus. Neuron, 92, 342–357.

Sheng, H. Z., Fields, R. D. & Nelson, P. G. 1993. Specific regulation of immediate early genes by patterned neuronal activity. J Neurosci Res, 35, 459–67.

Sheng, M. & Greenberg, M. E. 1990. The regulation and function of c-fos and other immediate early genes in the nervous system. Neuron, 4, 477–85.

Sherrin, T., Blank, T., Hippel, C., Rayner, M., Davis, R. J. & Todorovic, C. 2010. Hippocampal c-Jun-N-terminal kinases serve as negative regulators of associative learning. J Neurosci, 30, 13348–61.

Shi, R., Redman, P., Ghose, D., Hwang, H., Liu, Y., Ren, X., Ding, L. J., Liu, M., Jones, K. J. & Xu, W. 2017. Shank Proteins Differentially Regulate Synaptic Transmission. ENEURO, 4, Eneuro.0163-15.2017.

Shpokayte, M., Mckissick, O., Guan, X., Yuan, B., Rahsepar, B., Fernandez, F. R., Ruesch, E., Grella, S. L., White, J. A., Liu, X. S. & Ramirez, S. 2022. Hippocampal cells segregate positive and negative engrams. Commun Biol, 5, 1009.

Sloviter, R. S. & Lomo, T. 2012. Updating the lamellar hypothesis of hippocampal organization. Front Neural Circuits, 6, 102.

Stahl, P. L., Salmen, F., Vickovic, S., Lundmark, A., Navarro, J. F., Magnusson, J., Giacomello, S., Asp, M., Westholm, J. O., Huss, M., Mollbrink, A., Linnarsson, S., Codeluppi, S., Borg, A., Ponten, F., Costea, P. I., Sahlen, P., Mulder, J., Bergmann, O., Lundeberg, J. & Frisen, J. 2016. Visualization and analysis of gene expression in tissue sections by spatial transcriptomics. Science, 353, 78–82.

Stanfield, B. B. & Cowan, W. M. 1979. The morphology of the hippocampus and dentate gyrus in normal and reeler mice. J Comp Neurol, 185, 393–422.

Stuchlík, A., Petrásek, T., Prokopová, I., Holubová, K., Hatalová, H., Valeš, K., Kubík, Š., Dockery, C. & Wesierska, M. 2013. Place Avoidance Tasks as Tools in the Behavioral Neuroscience of Learning and Memory. Physiological Research, S1–S19.

Sullivan, K. E., Kendrick, R. M. & Cembrowski, M. S. 2021. Elucidating memory in the brain via single-cell transcriptomics. J Neurochem, 157, 982–992.

Sun, X., Bernstein, M. J., Meng, M., Rao, S., Sorensen, A. T., Yao, L., Zhang, X., Anikeeva, P. O. & Lin, Y. 2020. Functionally Distinct Neuronal Ensembles within the Memory Engram. Cell, 181, 410–423 e17.

Sun, X. & Lin, Y. 2016. Npas4: Linking Neuronal Activity to Memory. Trends Neurosci, 39, 264–275.

Sweis, B. M., Mau, W., Rabinowitz, S. & Cai, D. J. 2021. Dynamic and heterogeneous neural ensembles contribute to a memory engram. Curr Opin Neurobiol, 67, 199–206.

Takeuchi, T., Duszkiewicz, A. J. & Morris, R. G. 2014. The synaptic plasticity and memory hypothesis: encoding, storage and persistence. Philos Trans R Soc Lond B Biol Sci, 369, 20130288.

Terranova, J. I., Yokose, J., Osanai, H., Marks, W. D., Yamamoto, J., Ogawa, S. K. & Kitamura, T. 2022. Hippocampal-amygdala memory circuits govern experience-dependent observational fear. Neuron, 110, 1416–1431 e13.

Terranova, J. I., Yokose, J., Osanai, H., Ogawa, S. K. & Kitamura, T. 2023. Systems consolidation induces multiple memory engrams for a flexible recall strategy in observational fear memory in male mice. Nat Commun, 14, 3976.

The Gene Ontology, C. 2019. The Gene Ontology Resource: 20 years and still GOing strong. Nucleic Acids Res, 47, D330–D338.

Tonegawa, S., Morrissey, M. D. & Kitamura, T. 2018. The role of engram cells in the systems consolidation of memory. Nat Rev Neurosci, 19, 485–498.

Tonegawa, S., Pignatelli, M., Roy, D. S. & Ryan, T. J. 2015. Memory engram storage and retrieval. Curr Opin Neurobiol, 35, 101–9.

Tyssowski, K. M., Destefino, N. R., Cho, J. H., Dunn, C. J., Poston, R. G., Carty, C. E., Jones, R. D., Chang, S. M., Romeo, P., Wurzelmann, M. K., Ward, J. M., Andermann, M. L., Saha, R. N., Dudek, S. M. & Gray, J. M. 2018. Different Neuronal Activity Patterns Induce Different Gene Expression Programs. Neuron, 98, 530–546 e11.

Van Strien, N. M., Cappaert, N. L. & Witter, M. P. 2009. The anatomy of memory: an interactive overview of the parahippocampal-hippocampal network. Nat Rev Neurosci, 10, 272–82.

Vanrobaeys, Y., Mukherjee, U., Langmack, L., Beyer, S. E., Bahl, E., Lin, L. C., Michaelson, J. J., Abel, T. & Chatterjee, S. 2023. Mapping the spatial transcriptomic signature of the hippocampus during memory consolidation. Nat Commun, 14, 6100.

Wachi, T., Cornell, B. & Toyo-Oka, K. 2017. Complete ablation of the 14-3-3epsilon protein results in multiple defects in neuropsychiatric behaviors. Behav Brain Res, 319, 31–36.

Wang, M., Zhao, Y. & Zhang, B. 2015. Efficient Test and Visualization of Multi-Set Intersections. Scientific Reports, 5, 16923.

Weeber, E. J., Atkins, C. M., Selcher, J. C., Varga, A. W., Mirnikjoo, B., Paylor, R., Leitges, M. & Sweatt, J. D. 2000. A Role for the β Isoform of Protein Kinase C in Fear Conditioning. The Journal of Neuroscience, 20, 5906–5914.

Wesierska, M., Dockery, C. & Fenton, A. A. 2005. Beyond memory, navigation, and inhibition: behavioral evidence for hippocampus-dependent cognitive coordination in the rat. J Neurosci, 25, 2413–9.

Wu, T., Hu, E., Xu, S., Chen, M., Guo, P., Dai, Z., Feng, T., Zhou, L., Tang, W., Zhan, L., Fu, X., Liu, S., Bo, X. & Yu, G. 2021. clusterProfiler 4.0: A universal enrichment tool for interpreting omics data. Innovation (Camb), 2, 100141.

Yao, Z., Van Velthoven, C. T. J., Nguyen, T. N., Goldy, J., Sedeno-Cortes, A. E., Baftizadeh, F., Bertagnolli, D., Casper, T., Chiang, M., Crichton, K., Ding, S. L., Fong, O., Garren, E., Glandon, A., Gouwens, N. W., Gray, J., Graybuck, L. T., Hawrylycz, M. J., Hirschstein, D., Kroll, M., Lathia, K., Lee, C., Levi, B., Mcmillen, D., Mok, S., Pham, T., Ren, Q., Rimorin, C., Shapovalova, N., Sulc, J., Sunkin, S. M., Tieu, M., Torkelson, A., Tung, H., Ward, K., Dee, N., Smith, K. A., Tasic, B. & Zeng, H. 2021. A taxonomy of transcriptomic cell types across the isocortex and hippocampal formation. Cell, 184, 3222–3241 e26.

Yap, E. L. & Greenberg, M. E. 2018. Activity-Regulated Transcription: Bridging the Gap between Neural Activity and Behavior. Neuron, 100, 330–348.

Zovkic, I. B., Paulukaitis, B. S., Day, J. J., Etikala, D. M. & Sweatt, J. D. 2014. Histone H2A.Z subunit exchange controls consolidation of recent and remote memory. Nature, 515, 582–6.

