## Supplemental Figures and Legends for "Spatially Resolved Transcriptomic Signatures of Hippocampal Subregions and *Arc*-Expressing Ensembles in Active Place Avoidance Memory"

### 1 SUPPLEMENTAL FIGURES

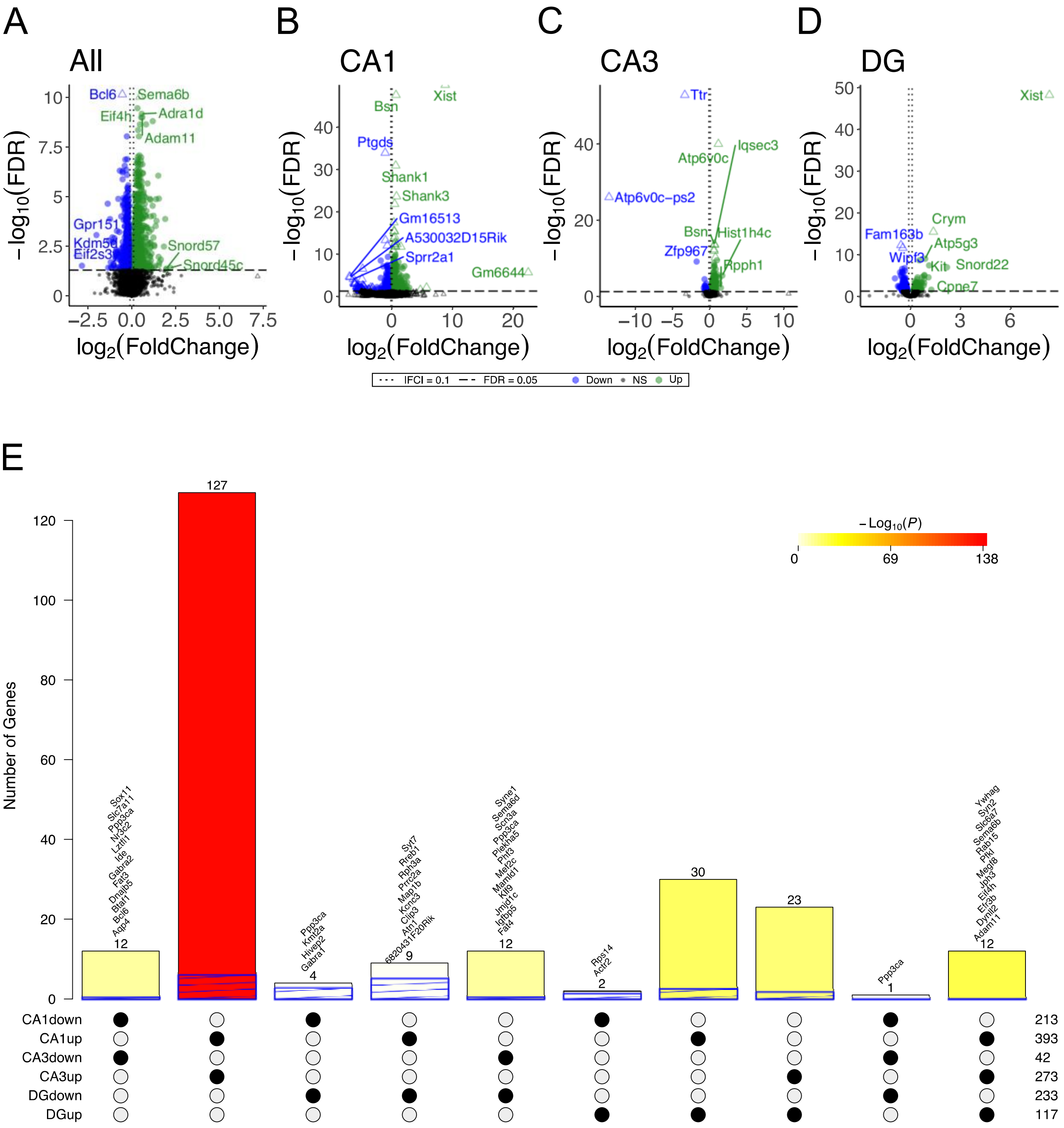


Supplemental Figure 1. Trained CA1 and CA3 subregions express similar DEGs and postsynaptic cellular component GO terms. **(A-D)** Volcano plot of DEGs between trained and untrained CA1, CA3 and DG samples, combined and individually. Triangular points mark data out of the frame of the main text figure. **(E)** Upset plot of overlaps in regional DEGs demonstrates greatest similarity between CA1 and CA3 samples. DEGs from each regional analysis were stratified by direction of fold change. In the upset plot, each bar represents the number of genes in an intersection of lists of DEGs. The intersecting lists is represented by the dots underneath the bar. The hashed line on each bar represents the expected intersection of two identically sized random lists. Overlaps were tested with Fisher's exact test using the list of all expressed genes as a background. The color of the bar represents the $-\log_{10}(P)$ for the BH corrected P-value of each intersection.

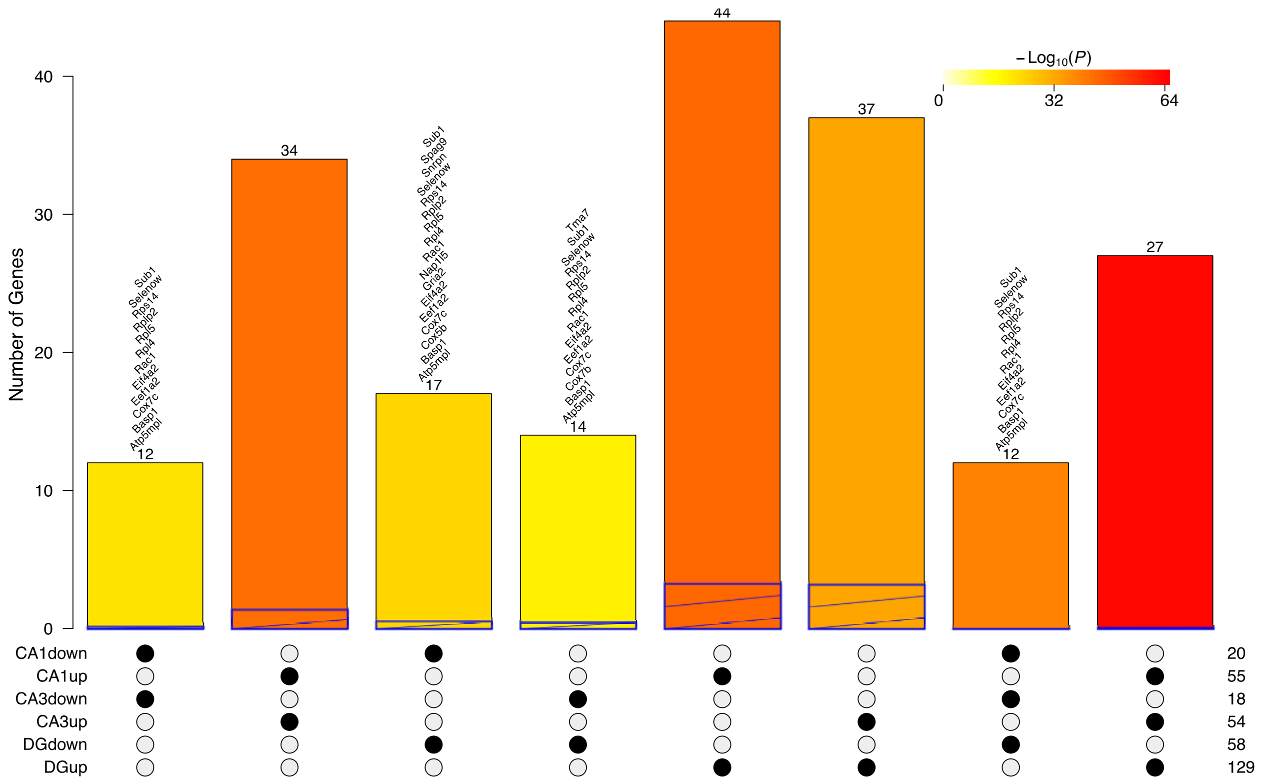


Supplemental Figure 2. Upset plot of overlaps in regional DEGs demonstrates greatest similarity between CA1 and DG subregions. DEGs from each regional analysis were stratified by direction of fold change. In the upset plot, each bar represents the number of genes in an intersection of lists of DEGs. The intersecting lists is represented by the dots underneath the bar. The hashed line on each bar represents the expected intersection of two identically sized random lists. Overlaps were tested with Fisher's exact test using the list of all expressed genes as a background. The color of the bar represents the $-\log_{10}(P)$ for the BH corrected P-value of each intersection.

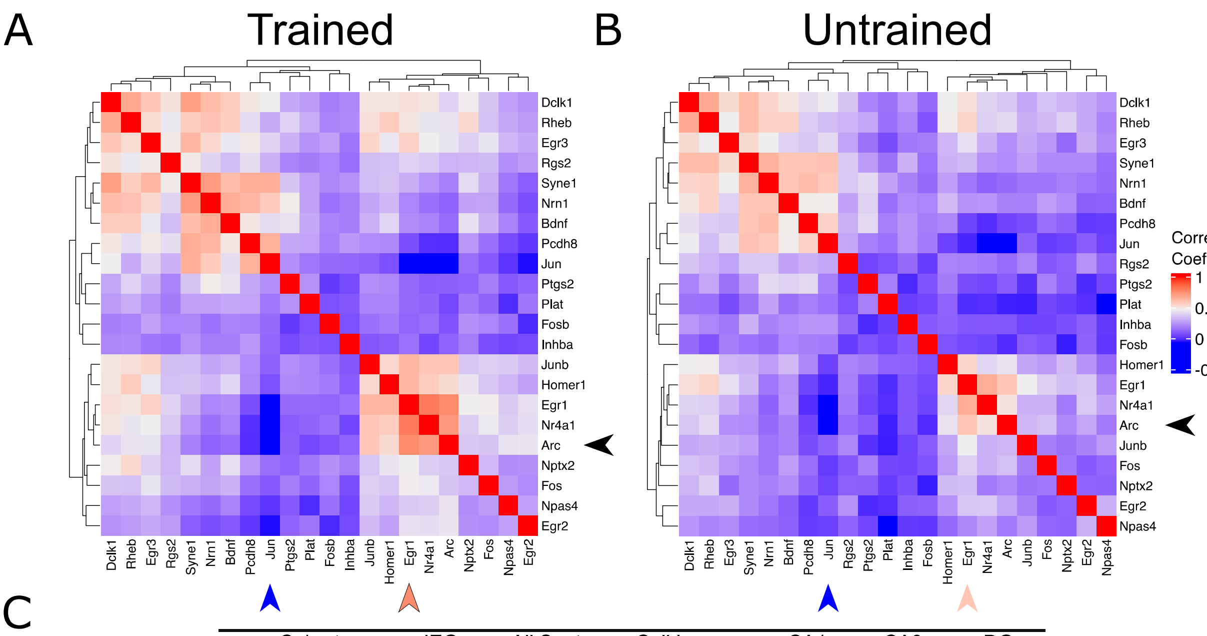


C

| Cohort | IEG | All Spots | Cell Layers | CA1 | CA3 | DG |
|---|---|---|---|---|---|---|
| Trained | Arc | 195/542 | 56/146 | 34/49 | 20/43 | 2/54 |
| | Egr1 | 380/542 | 130/146 | 48/49 | 41/43 | 41/54 |
| | c-Jun | 308/542 | 106/146 | 22/49 | 32/43 | 52/54 |
| Untrained | Arc | 169/568 | 58/155 | 34/51 | 19/42 | 5/62 |
| | Egr1 | 256/568 | 113/155 | 51/51 | 33/42 | 29/62 |
| | c-Jun | 175/568 | 83/155 | 12/51 | 18/42 | 53/62 |

D

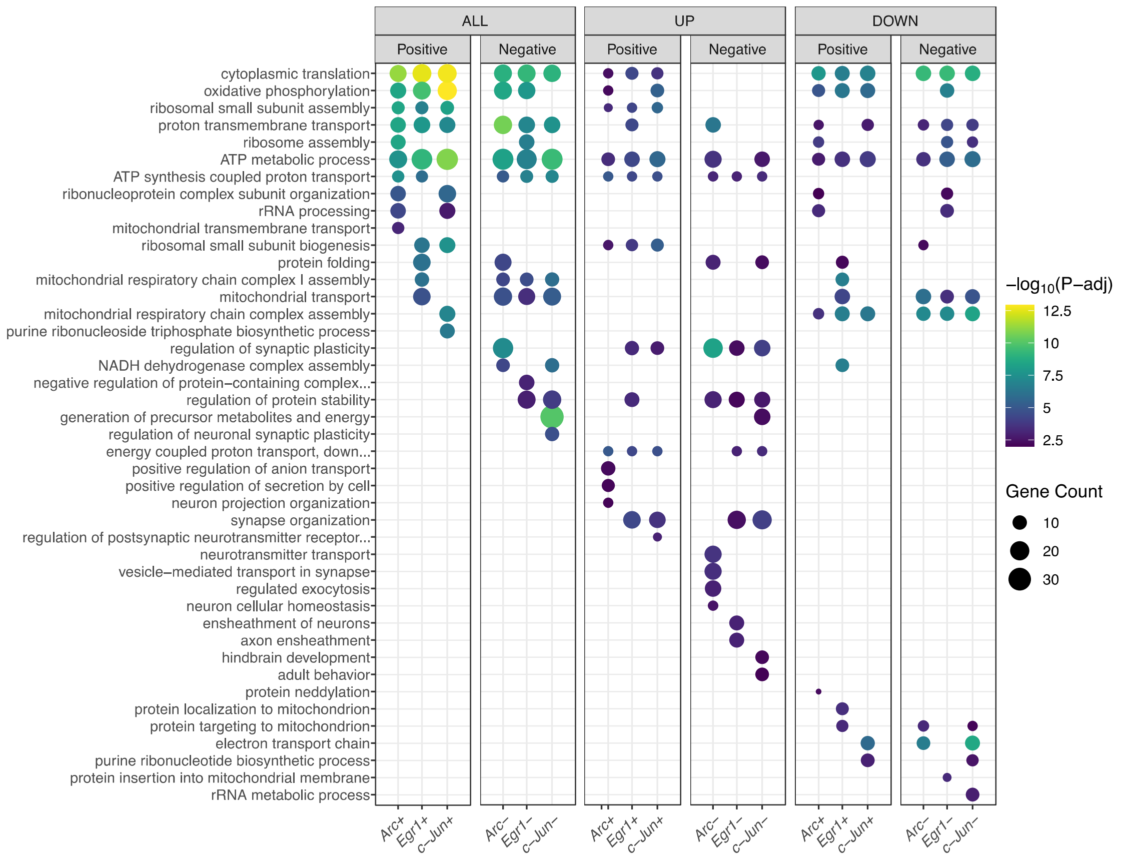

Supplemental Figure 3. The expression of the IEGs *Egr1* and *c-Jun* are most and least correlated with the expression of *Arc*, respectively. **(A, B)** Heatmaps reflecting the pairwise Pearson correlation coefficients between IEGs detected in the Trained **(A)** and Untrained **(B)** sample. Pearson correlation between *Arc* and *Egr1* or *c-Jun* are marked by the caret on the x-axis colored to reflect the value of the correlation coefficient. The correlation coefficients for *Arc* and *Egr1* are r(21) = .598, p = 2.24 x $10^{-56}$ in the untrained sample and r(21) = .764, p = 5.45 x $10^{-105}$ in the trained sample. The correlation coefficients for *Arc* and *c-Jun* are r(21) = -.057, p = 0.17 in the untrained sample and r(21) = -.105, p = 0.014 in the trained sample. **(C)** Table total number of IEG expressing spots relative to the total number of hippocampal spots for each area. **(D)** Biological processes overrepresented in the analysis of IEG-expressing spots across trained and untrained samples amongst all DEGs (left) and stratified by up- (middle) and down-regulated (right) DEGs. Dot color reflects the statistical significance ($-\log_{10}$(FDR)) of the biological process enrichment. Dot size reflects the number of detected DEGs mapping to the gene belonging to a given biological process.

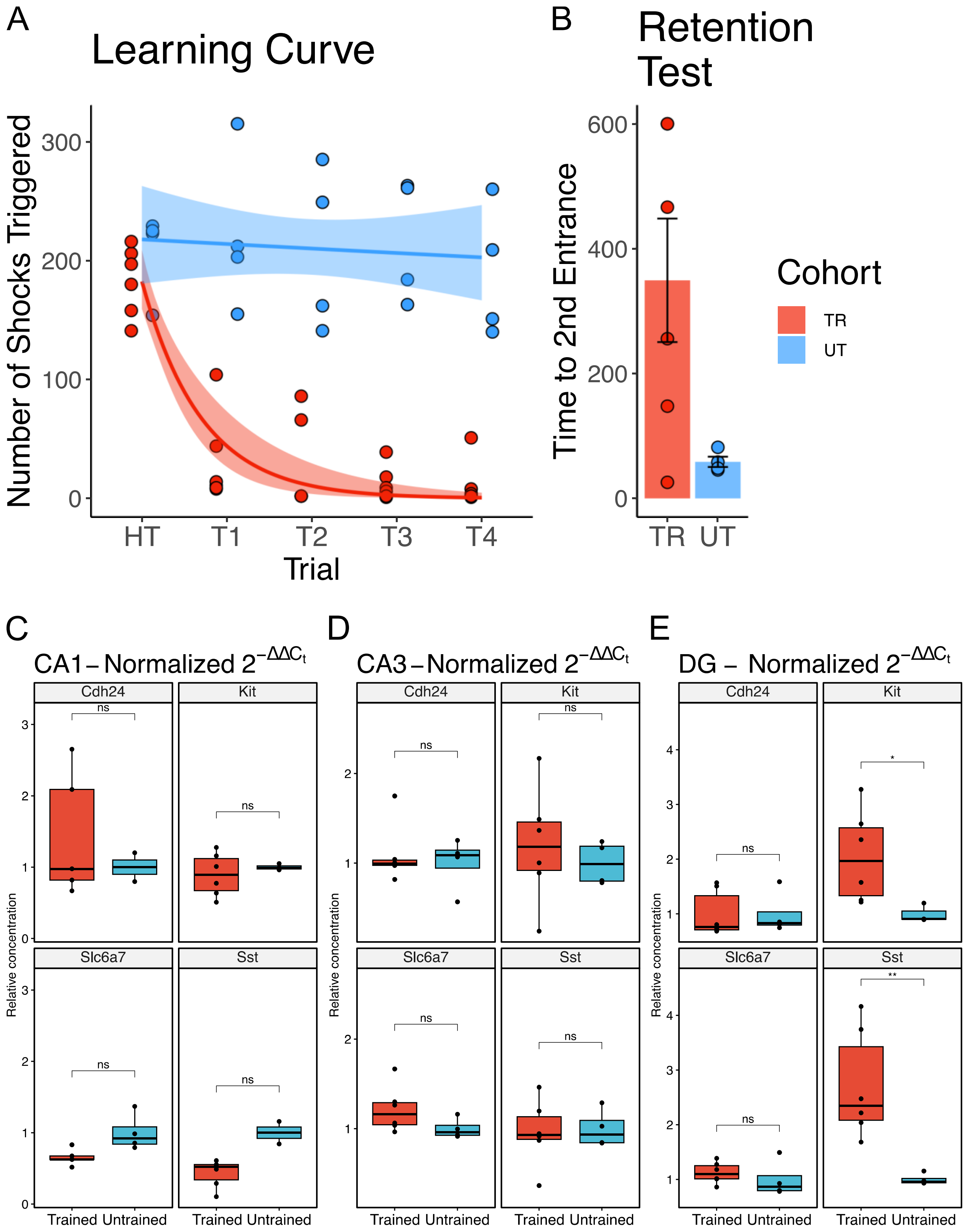
A
Learning Curve
Number of Shocks Triggered
300
200
100
0
HT
T1
T2
T3
T4
Trial
B
Retention Test
Time to 2nd Entrance
600
400
200
0
TR
UT
Cohort
TR
UT
C
CA1 – Normalized $2^{-\Delta\Delta C_t}$
D
CA3 – Normalized $2^{-\Delta\Delta C_t}$
E
DG – Normalized $2^{-\Delta\Delta C_t}$
Relative concentration
Cdh24
Kit
Slc6a7
Sst
ns
*
**
Trained Untrained

38

Supplemental Figure 4. Relative expression of select DEGs through RT-qPCR. **(A,B)** RT-qPCR was performed on samples collected from animals trained under conditions identical to those used for bulk RNA sequencing or spatial transcriptomics. **(A)** Training performance measured as number of shocks triggered. Trained mice learned to avoid the location of the shock zone indicated by the decreased number of shocked triggered. Exponential fit learning curves included with shaded error bars (SE). **(B)** Memory performance measured as time to second entrance in the retention test trial. Trained mice demonstrated higher time to second entrance during the retention test. Trained mean 349s ± 98.9 (SE), Untrained mean 58.5s ± 8.23 (SE). **(C-E)** Average normalized relative expression ($2^{-\Delta\Delta Ct}$) of genes in trained samples (red) as compared to untrained samples (blue) in CA1 **(C)**, CA3 **(D),** and DG **(E)** subregions. n= 6 trained and n=4 untrained animals. Student t-test was performed between $\Delta$Ct values from trained and untrained samples for each gene.
